# A pan-cancer catalog of mutagenesis at transcription factor binding sites

**DOI:** 10.64898/2026.10.19.760246

**Authors:** Harshit Sahay, Yuncheng Duan, Raluca Gordân

## Abstract

Transcription factor (TF) binding can contribute to mutagenesis by altering DNA damage formation or interfering with DNA repair. Although these effects have been characterized for selected TFs and mutagenic processes, their prevalence across TFs and cancer types remains poorly understood. A systematic characterization of mutation enrichment at TF binding sites is needed to guide mechanistic studies and improve the interpretation of noncoding mutations in cancer.

Here, we analyzed somatic mutations from over 4,000 tumors across 15 tissue types to generate a pan-cancer catalog of mutation patterns at binding sites for 655 TFs, organized into 276 motif clusters. We assessed mutation enrichment at single-nucleotide resolution, accounting for sequence-dependent and local variation in background mutation rates. Nearly one third of <TF-cluster, tissue> pairs with sufficient mutation counts showed significant enrichment at one or more binding site positions. Enrichment was most widespread in melanoma but was detected in all 15 tissue types, with the affected TFs and binding site positions varying across tissues. Highly expressed TFs were more likely to show enrichment than lowly expressed TFs, consistent with a role for TF binding in increased mutagenesis at the binding sites. Mutational signature analysis implicated processes beyond UV damage, including age-associated processes, deamination lesions, oxidative damage, and tobacco exposure, as potentially contributing to TF-induced mutagenesis. Together, these findings suggest that diverse interactions between TF binding, DNA damage, and DNA repair may contribute to mutation enrichment in regulatory DNA.

Our pan-cancer catalog reveals position-specific enrichment of somatic mutations at TF binding sites across all 15 tissue types analyzed. By identifying specific TFs and binding site positions enriched for somatic mutations, together with potential mutagenic processes acting at these positions, our study provides a roadmap for experimental assessment of TF-induced mutagenesis and informs the interpretation and prioritization of non-coding mutations in cancer.

## INTRODUCTION

The advent of whole-genome sequencing has recently provided detailed maps of somatic mutations for thousands of human tumors (The ICGC/TCGA Pan-Cancer Analysis of Whole Genomes Consortium 2020). Analyses of these data revealed a striking level of variation in somatic mutation rates along the genome and provided new insights into mutagenic mechanisms acting at different scales (Hodgkinson et al. 2012; Lawrence et al. 2013; Supek and Lehner 2019). At the megabase scale, the variation in mutation rates has been tied to differences in chromatin organization, replication timing, and DNA repair activity (e.g. (Stamatoyannopoulos et al. 2009; Schuster-Bockler and Lehner 2012; Polak et al. 2015; Supek and Lehner 2015)). At the kilobase scale, the mutation rate is lower at accessible regulatory regions than in their neighboring flanks, potentially due to open chromatin providing easier access to DNA repair machineries (Polak et al. 2014). However, within these open chromatin regions, binding sites of transcription factor (TF) proteins are oftentimes hypermutated (Katainen et al. 2015; Perera et al. 2016; Sabarinathan et al. 2016). This is not explained by selection alone (Kaiser et al. 2016), leading to the hypothesis that TF binding is potentially mutagenic (Reijns et al. 2015; Sabarinathan et al. 2016).

TF-induced mutagenesis has important implications for identifying driver regulatory mutations that contribute to tumorigenesis. TF binding to target genomic sites drives gene regulation, disruption of which plays a major role in tumor growth (Hodgkinson et al. 2012; Huang et al. 2013; Hnisz et al. 2016). Mutations in non-coding DNA account for >90% of all somatic mutations (Khurana et al. 2016) and oftentimes fall within regulatory elements bound by TFs. Identifying regulatory drivers within non-coding DNA remains challenging even when a large number of tumors are analyzed (Cuykendall et al. 2017; Rheinbay et al. 2020; Elliott and Larsson 2021). A widespread pattern of hypermutation in TF binding sites, driven by TF binding as opposed to selection in the cancer cells, can significantly hinder efforts to identify candidate regulatory drivers. To assist in these efforts, in our current study we comprehensively catalog the hypermutation patterns at TF binding sites across a wide range of TFs and tumor types, at single-nucleotide resolution, and we generate testable hypotheses about the putative mechanisms leading to these mutation patterns, as a key first step toward a robust understanding of mutagenic processes in regulatory DNA. Understanding and characterizing these processes is critical for building accurate models of background mutation rates in regulatory DNA, which in turn will help in future efforts to identify regulatory driver mutations.

The pattern of hypermutation at TF binding sites is widespread, but not identical across factors and tumor types. For example, binding sites of CTCF show an enrichment of mutations in skin, colorectal, and gastric tumors (Katainen et al. 2015; Poulos et al. 2016; Umer et al. 2016). Still, the magnitude of this enrichment and the precise positions within CTCF binding sites that show an elevated mutation rate are distinct among tumor types (Pinoli et al. 2020). In melanoma, many TFs show enrichment of somatic mutations in their binding sites (Sabarinathan et al. 2016), but the magnitude of the enrichment is variable across TF families (Frigola et al. 2021). In prostate tumors, it has been reported that genomic regions bound by AR, but not other TFs, are enriched for somatic mutations (Morova et al. 2020). These prior observations suggest that the role of TF binding in mutagenesis is likely specific to the tumor type and the identity of the TF in question, thereby prompting us to perform a comprehensive characterization of the mutation enrichment patterns in TF binding sites over a wide range of tumors and factors.

The molecular mechanisms leading to the enrichment of somatic mutations in TF binding sites are also likely to be TF- and tumor type-specific. Some studies attribute this enrichment to the inhibition of DNA repair pathways (Sabarinathan et al. 2016; Frigola et al. 2021), a hypothesis consistent with genome-wide maps of DNA repair activity in human and yeast cells (Adar et al. 2016; Hu et al. 2017; Duan et al. 2022). Under this hypothesis, TF inhibition of DNA repair is proposed to occur as a result of TF binding to DNA lesions, thereby preventing recognition of the lesions by DNA repair enzymes. However, this requires TFs to efficiently bind to damaged DNA substrates, and little is known about how DNA damage affects TF binding. We recently showed that TFs can bind to DNA mismatches (Afek et al. 2020) and to UV lesions (Mielko et al. 2023) with binding strengths similar to or even higher than canonical Watson-Crick binding sites, making the proposed mechanism of TF-repair competition plausible. However, each TF has its own pattern of binding to damaged DNA sites, and further experiments are needed to understand these interactions and eventually build predictive models of TF binding to damaged DNA and TF- repair competition. A second mechanism potentially responsible for the enrichment of mutations at TF binding sites is an increase in DNA damage formation, likely due to DNA conformational changes induced by TF binding (Tornaletti and Pfeifer 1995; Elliott et al. 2018; Mao et al. 2018; Frigola et al. 2021; Wasserman et al. 2026). Given that different TFs induce different distortions upon DNA binding, any effects on damage formation, and subsequently on mutation rates, are expected to be specific to each TF and damage type.

Overall, the diverse mechanisms that potentially lead to the enrichment of somatic mutations in TF binding sites, together with the variation in enrichment among TFs and tumor types, make experimental validation very difficult. Since TF binding, DNA damage, and DNA repair mechanisms are all sequence-specific, any experimental efforts to characterize the interplay between TF binding and DNA damage and repair require precise hypotheses, ideally at single-base pair resolution. Toward this goal, in the current study we built a high-resolution, pan-cancer catalog of somatic mutation patterns at TF binding sites. We analyze these patterns to identify binding site positions with a statistically significant enrichment of somatic mutations, and link them to the DNA damage and repair pathways likely to be involved. Our work provides am important map for future experimental work to test hypotheses of TF-induced mutagenesis, by nominating specific TFs and mutagenic mechanisms to be tested.

## RESULTS

Previous studies of mutation enrichment at TF binding sites focused on a small number of specific TFs (Morova et al. 2020), or on specific cancer types (Sabarinathan et al. 2016; Frigola et al. 2021). Other studies analyzed ∼100 TFs, but not at single-base pair resolution (Kaiser et al. 2016). To characterize the putative effects of TF binding on mutagenesis more systematically, we first sought to build a pan-cancer catalog of mutation rates in binding sites compared to their flanking regions, and to identify TFs for which the mutation rates within the binding sites are significantly higher than expected by chance. Toward this goal, we collected publicly available somatic mutation data from the International Cancer Genome Consortium (ICGC) (Zhang et al. 2019) and TF-DNA binding motif models for 655 human TFs organized in 276 TF motif clusters (Vierstra et al. 2020). Our analysis comprised of 69,767,173 somatic mutations from 36 ICGC projects across 15 primary tissues, representing the most comprehensive evaluation to date of mutagenesis at TF binding sites in cancer cells (**Table S1**).

### TF binding sites show elevated mutation rates across a wide range of cancer types

To calculate mutation rates at TF binding sites, we first need to identify putative TF-bound sites across the human genome in each cancer type/tissue of interest. For this purpose, we focused on open chromatin regions that contained TF-DNA motif matches for all motifs compiled in (Vierstra et al. 2020). Regions of open chromatin were defined using DNase-seq and ATAC-seq data sets from both primary tissues and tumor cells (Thurman et al. 2012; Roadmap Epigenomics et al. 2015; Corces et al. 2018), as detailed in **Table S1** and **Methods**.

Mutation rates in accessible TF binding sites are likely to show tissue-specific patterns, as TF binding and DNA accessibility are different among tissue types. As a result, we performed our analysis of mutational patterns at TF binding sites by considering each tissue type separately. Within each tissue of interest, different tumor types have both shared and unique mutagenic processes acting on the genome. While these tumor types belonging to the same tissue would ideally be analyzed separately, the small number of mutations per specific tumor type, when further intersected with TF binding site locations in accessible chromatin, make it difficult to identify statistically significant patterns. To address this problem, when considering each of the 15 tissue types of interest, we aggregated mutations observed in all ICGC projects with tumor types relevant to that tissue (**Table S1**), which improves our ability to identify TFs that show hypermutation patterns in their binding sites. As more mutation data becomes available, our analysis can be refined for tumor types within each tissue, to uncover further nuances of mutational patterns in these binding sites.

The collection of TF-DNA binding motifs compiled and clustered by Vierstra et al. was used to call, for each motif cluster, putative TF binding sites across the human genome, as in their original study (Vierstra et al. 2020). Next, we intersected these TF binding site calls with the open chromatin regions for each tissue type, and we aggregated tissue-specific somatic mutations within the binding sites and their flanking regions for each TF motif cluster. We note that a cluster may correspond to multiple paralogous TFs with similar DNA binding specificities and thus similar DNA motifs, for which TF binding site calls cannot be distinguished without additional data (such as ChIP-seq, which is not available on a large scale for the tumor/tissue types and TFs considered).

Our aggregate analysis of somatic mutations across a wide range of TFs showed that TF binding sites have a clear elevation in mutation rates compared to their flanking regions (**Fig. 1a**), a trend observed for most tissue types tested (**Fig. 1b; Fig. S1a).** We further observed that this elevation was oftentimes localized at the sites of TF binding (**Fig. 1c**). For certain TFs, the elevation was also most pronounced for high-affinity binding sites (**Fig. 1d**, top panel), with moderate elevation seen at medium-affinity binding sites (**Fig. 1d**, middle panel), and no elevation seen at binding sites with low affinity (**Fig. 1d**, bottom panel). This correlation between the increase in mutation rate at binding sites (compared to their flanking regions) and the strength of TF binding suggested that TF binding, rather than some broader property of the bound regions, is potentially causing the observed hypermutation patterns (**Fig. 1a-d, Fig. S1**). Previous work elegantly combined genomic maps of DNA damage and repair to implicate both higher damage formation and reduced repair activity at TF binding sites as potential mechanisms for the increased mutation rates at these sites (Sabarinathan et al. 2016; Frigola et al. 2021; Wasserman et al. 2026). Beyond such correlation-based analyses, however, there is limited direct experimental evidence (Mao et al. 2018; Zhu et al. 2025) in support of these mechanisms. This is not surprising given the large variety of TFs and mutagenic processes that need to be considered for testing hypotheses of TF-induced mutagenesis at binding sites. The analysis framework developed here can provide clues regarding potential mechanisms for each TF and tumor/tissue type, which can then be followed up experimentally.

**Figure 1:**
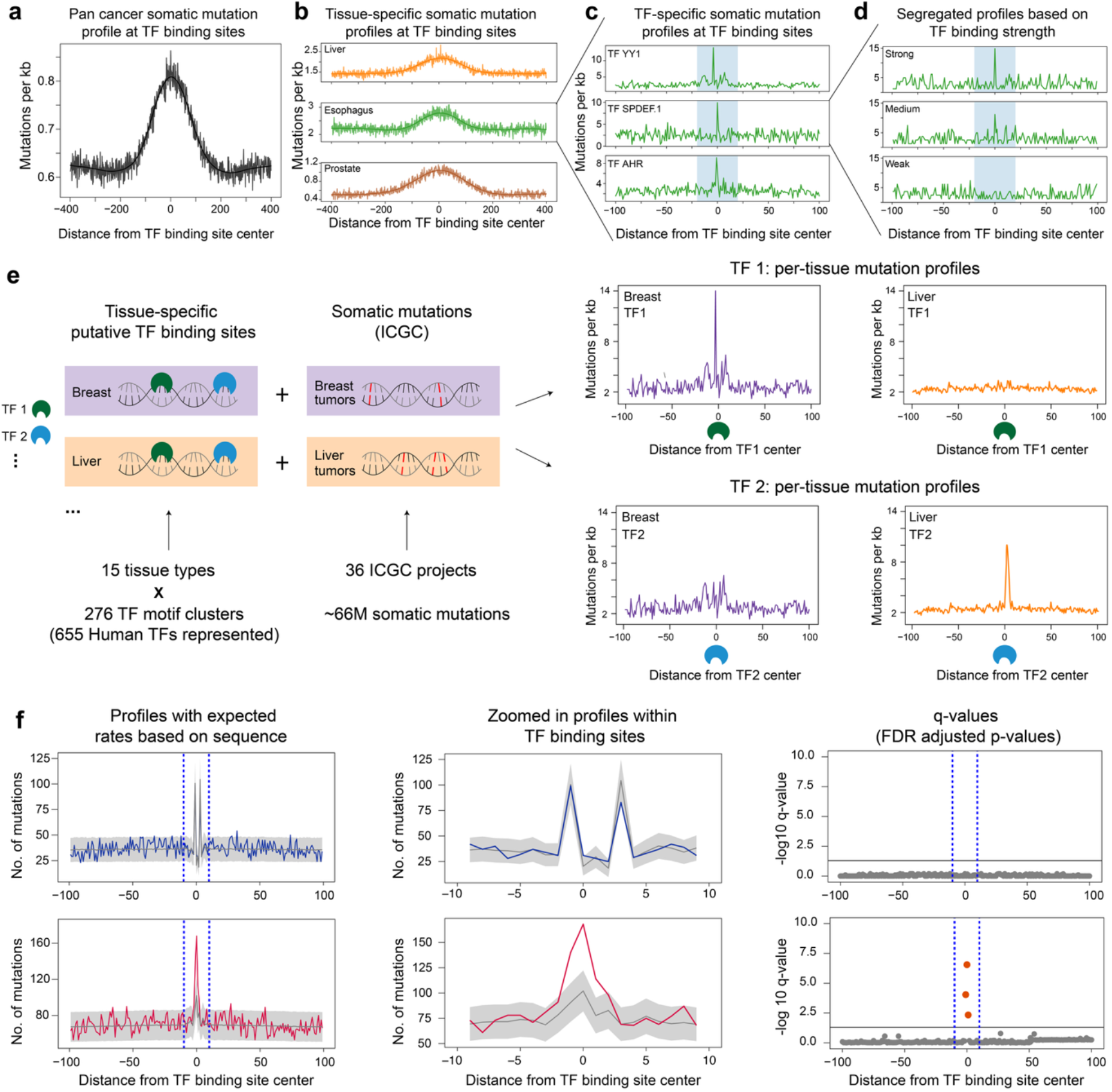
Pan-cancer analysis of hypermutation patterns at transcription factor (TF) binding sites. **(a)** TF binding sites in open chromatin show an elevation in somatic mutation counts compared to their flanking regions. The mutation counts displayed were computed by aggregating binding sites for all TFs and mutations from all tumors analyzed (**Methods**). **(b)** Tumors in several tissue types show an elevation of somatic mutations at TF binding sites. Liver (orange), esophagus (green), and prostate (brown) are shown here. **(c)** Examples of TFs with mutation peaks at their binding sites (here, in esophageal tumors). **(d)** The association between a TF’s binding strength and the observed mutation count elevation is suggestive of a role for TF binding in mutagenesis. Profiled here is TF cluster SPDEF/1 in esophageal tumors. ‘Strong’, ‘Medium’, and ‘Weak’ sites were obtained by separating all binding sites used in panel c into 3 quantiles based on motif scores (**Methods**). **(e)** For each of the 276 TF clusters, we performed mutational analyses in a tissue-specific manner, using tissue-specific open chromatin regions along with somatic mutations from relevant tumors (**Methods**). Shown here are examples for two such TFs, TF 1 (green) and TF 2 (blue), in two tissue types (purple: breast, orange: liver). For this set, our analysis yields 4 mutation profiles, one for each <TF-cluster, tissue> pair. In this example, a local hypermutation pattern is seen at TF 1 binding sites in breast tumors and at TF 2 binding sites in liver tumors, but not in other <TF-cluster, tissue> combinations. **(f)** Local hypermutation at TF binding sites can sometimes be explained by sequence alone. To identify and filter out such cases, we augment our mutation profiles with background mutation counts calculated from trinucleotide mutation rates for each tumor (**Methods**). Shown here are two mutation profiles (left panel), along with the mean +/- 2 standard deviations of the sequence-derived expected mutation counts (gray). Mutation profiles for the two TFs within the binding site (center panel) illustrate the difference between cases where the mutation elevation within binding sites is explained by sequence (blue) versus cases where it is not (red). The latter is more likely to result from binding of the TF. We develop a workflow to systematically identify positions within the binding site where such deviations from the background sequence model are seen (**Methods**). This results in a position-specific p-value (right), that can be used to identify all positions where TF binding could be implicated in mutagenesis (red dots).

Since TF binding and mutagenic mechanisms are both tissue-specific, the mutation rates at TF binding sites are expected to differ in different tissue contexts. Indeed, when analyzing the overall elevation in mutation rates at TF binding sites compared to flanking regions, we found that the magnitude of this elevation (**Methods**) differs among tumor/tissue types, from 90% in prostate cancers to 7% in brain cancers (**Fig. S1a**). Different tumor/tissue types also showed variation in the number of TFs and TF clusters that showed an elevation in mutation rates at the binding site, with skin tumors showing an elevation for 153 (out of 276) clusters, encompassing 351 TF proteins, compared to 31 clusters encompassing 107 TFs for stomach tumors (**Fig. S1b**). The precise number of TFs and TF clusters for which elevated mutation rates can be reliably detected depends on the total number of mutations available for each tissue type (**Fig. S1a, Methods**). When considering the set of TF clusters with sufficient mutation counts to perform this analysis, between 20 and 65% of TFs showed an elevation in the different tissue types, suggesting that the pan-TF trends observed in **Fig. 1a** are widespread across individual TFs.

When focusing on individual TFs, we also found that their binding site mutation profiles differ substantially among tissue types (**Fig. S1c**), suggesting that each <TF, tissue> pair should be analyzed separately. To account for this, we identified putative binding sites for all 276 TF clusters within the open chromatin regions of each tissue separately (**Methods**). These maps of TF binding were then combined with the somatic mutations across all tumors in that tissue to produce one mutation profile for every <TF-cluster, tissue> pair (**Fig. 1e**). Our analysis thus enables the systematic identification of specific TFs that show binding site mutation counts significantly higher than their immediate flanking regions, and thus could be involved in mutagenesis in particular tumors/tissues, as described below.

### Elevated mutation rates within TF binding sites are not explained by sequence

Since TFs bind to genomic sites with specific DNA sequence patterns, the local elevation of mutation rates at TF binding sites (**Fig. S1a,b**) could result simply from the fact that the underlying sequence follows a common pattern that matches the pattern of the mutagenic processes in that particular tumor/tissue (Yiu Chan et al. 2019; Liu et al. 2021). For example, transcription factor ETS1 binds with high affinity to sequences with a TTCC core motif, which contains three overlapping dipyrimidines (TT, TC, and CC). Since UV irradiation leads to the creation of photoproducts at dipyrimidines, genomic locations rich in TT, TC, CT, and CC dinucleotides are oftentimes hotspots of UV-induced DNA damage, as well as mutational hotspots in melanoma tumors (Pfeifer 2020). As a result, ETS1 binding sites, which contain these highly damageable dipyrimidines, are expected to show a higher mutation rate than their flanking regions, which have a more diverse sequence composition. This trend is indeed observed in analyses of mutation rates in melanomas (Sabarinathan et al. 2016; Mao et al. 2018; Wasserman et al. 2026). In order to accurately ascertain the impact of ETS1 binding on mutation rates, one must first determine to what extent the elevation observed at these binding sites can be explained by sequence alone. Previous work in melanomas showed that for ETS1, and some other TFs, the elevation patterns are not explained solely by sequence (Mao et al. 2018; Frigola et al. 2021). As a result, these TFs were proposed to play potentially mutagenic roles in melanoma tumors.

Here, we sought to comprehensibly determine cases where the elevation in mutation rates seen at TF binding sites is not entirely explained by sequence, across hundreds of TFs and more than a dozen tissue types. Doing so would enable us to determine when TF binding is likely to affect mutation rates, and thus nominate TFs that can play mutagenic roles in a given tumor.

To control for DNA sequence as a confounder when analyzing mutagenesis at TF binding sites, we modeled the overall mutation rate for each possible nucleotide in its immediate context, i.e. considering one nucleotide upstream and downstream, as in previous studies (Sabarinathan et al. 2016; Yang et al. 2018). Due to differences between open and closed chromatin (Polak et al. 2014), we filtered out inaccessible genomic regions and calculated mutation rates for each trinucleotide (defined as described above) within accessible DNA. In addition, to account for local variation in mutation rates due to differences in DNA replication timing, gene expression levels, or other mechanisms acting at large scales (Hodgkinson et al. 2012; Liu et al. 2013; Polak et al. 2015; Sherman et al. 2022), the trinucleotide mutation rates were corrected (**Methods, Fig. S2**) similarly to previous studies (Poulos et al. 2016; Sabarinathan et al. 2016).

Next, for every position within TF binding sites and their immediate flanks, we calculated an expected mutation count, and asked whether the observed mutation count at that position was significantly higher than expected (**Methods**). Briefly, this was done by modeling mutations across different binding sites as a Poisson-binomial process. At every position, mutation rates within the observed trinucleotides were modeled as independent Bernoulli processes that were then aggregated to obtain the expected distribution at that position. When TF binding does not contribute to mutagenesis, the observed mutation rate is expected to fall within the background distribution (**Fig. 1f**, top). However, if TF binding causes increased mutagenesis, one would expect the observed mutation count to be significantly higher than expected based on the background model (**Fig. 1f**, bottom). The Poisson-binomial process helps formalize this intuition by enabling us to calculate an expected mutation count, while the closed form (Hong 2013) allows us to calculate the p-value of observing any given mutation count under the null hypothesis of no effect of TF binding on mutagenesis (**Methods**).

Using this methodology, we designed a workflow to identify <TF-cluster, tissue> pairs that show significant enrichment for somatic mutations at the TF binding sites (**Fig. 1f**). Here, we defined a position within the binding sites of a particular TF as ‘enriched’ for somatic mutations in a tissue of interest if the mutation count at that binding site position, for that particular tissue, was significantly higher than the mutation count expected based on sequence alone. These are the positions where TF binding may contribute to the increased mutation rate, and thus should be the focus of future studies. Statistical significance was assessed based on p-values according to a Poisson-binomial process, as described above, adjusted for multiple hypothesis testing using the Benjamini-Hochberg correction (Benjamini and Hochberg 1995) applied to the TF binding sites and their flanking regions (+/-100 bp**; Methods**).

This strategy enabled us to identify binding site positions within <TF-cluster, tissue> pairs that not only showed an *elevation* of mutations compared to the flaking regions, but also an *enrichment* of mutations compared to the expected background mutation counts due to sequence alone (**Fig. 1f**). The observed enrichments are tissue-specific (**Fig. S1d**), as expected, underlining the importance of modeling these effects separately for each tissue. Our analysis thus suggests that the binding of certain TFs to DNA can cause more mutations than one would expect by chance, and this TF-induced mutagenesis mechanism may be more widespread than previously reported (Yang et al. 2018; Morova et al. 2020; Frigola et al. 2021).

### Widespread enrichment of somatic mutations at TF binding sites in cancer cells

To gain a broader understanding of the TFs and tissue types for which TF binding is likely to lead to increased mutagenesis at the binding sites, we applied our analysis workflow (**Fig. 1f**) to all 15 tissue types and 276 TF clusters analyzed in this study. Specifically, for each <TF-cluster, tissue> pair, we built mutation profiles as described above, with each profile augmented with expected mutation counts that control for sequence-based and local variation in mutation rates (**Fig. 1e-f, Methods**). This workflow yielded, for each position in a <TF-cluster, tissue> mutation profile: an observed somatic mutation count, a background distribution of expected mutation counts, and a q-value that reflects whether the deviation of the observed counts from the expected counts is statistically significant (**Fig. 1f**, **Methods**). For each <TF-cluster, tissue> pair, we performed the analysis described above for binding site calls in accessible DNA regions identified using both tumor-specific (ATAC-seq) and tissue/cell-line-specific (DNase-seq) accessibility datasets (**Methods**). The full results are available in **Data Sets 1-15**, for all 276 TF clusters and 15 tissue types analyzed.

To visualize the overall landscape of mutation enrichment at TF binding sites across a wide range of TFs and tissue types, for each <TF-cluster, tissue> pair we identified the most significant deviation (i.e. lowest q-value) observed within the binding site, defined as +/- 10 bp from the center of the TF motif. Positions with highly significant deviations (i.e. low q-values) are more likely to be implicated in TF-driven mutagenesis, since these are positions where TF binding is accompanied by a high deviation from the expected mutation counts (**Fig. 1f**, bottom). On the other hand, positions with large q-values either show mutation elevation that is explained by the underlying sequence of the binding sites (**Fig. 1f**, top), or no mutation elevation compared to flanking regions. Thus, our analysis allows us to differentiate between mutation *elevation* at TF binding sites (defined as any increase in mutations at binding sites relative to their flanking regions) versus mutation *enrichment* (defined as an increase in mutation counts beyond the range predicted by local sequence and other local covariates). Future experimental studies of TF-driven mutagenesis should focus on <TF-cluster, tissue> pairs that show an *enrichment* of mutations at one or more positions within the TF binding sites, rather than simply a pattern of mutation *elevation* in the binding site region. To help guide such future studies, we generated a comprehensive catalog of mutation enrichment at TF binding sites across all 15 tissue types analyzed here, and for all 276 TF clusters representing 655 human TFs (**Data Sets 1-15**). In addition, for each of the 4,140 (276x15) <TF-cluster, tissue> pairs, we report the minimum q-value across all positions in the TF binding sites (**Table S2**), and we visualize these results in **Fig. 2a**.

**Figure 2:**
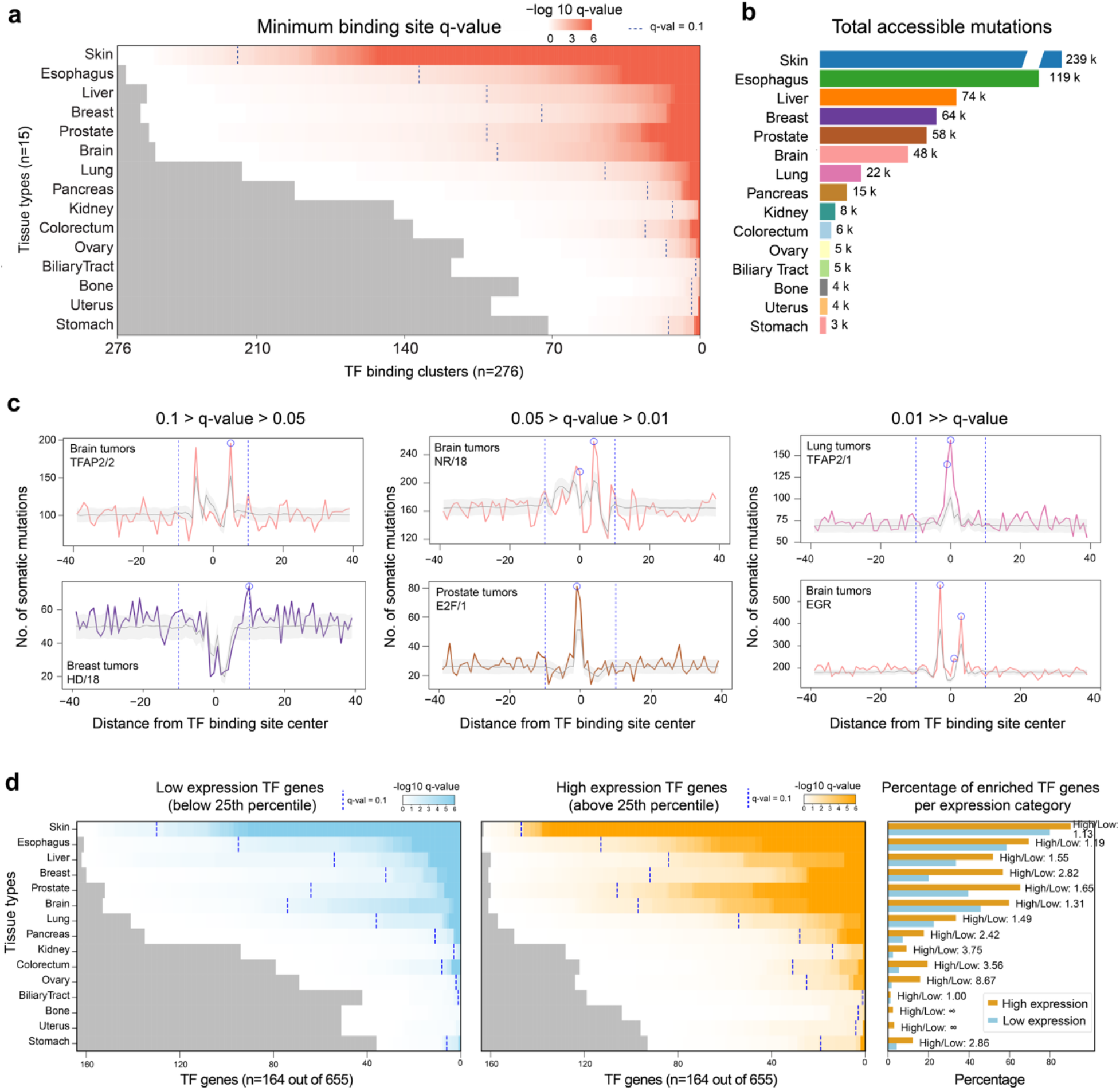
Enrichment of cancer somatic mutations at TF binding sites across tissue types. **(a)** Heatmap of the most significant enrichment, i.e. the largest -log10(q-value), observed for each <TF-cluster, tissue> pair (**Methods**). Tissue types were sorted, top to bottom, in decreasing order of the total number of mutations in accessible genomic regions. For each row (i.e. tissue type), the TF clusters were sorted in decreasing order of their most significant q-values (calculated across all binding site positions for that particular <TF-cluster, tissue> pair). Vertical blue lines indicate the q-value cutoff of 0.1. Gray rectangles mark <TF-cluster, tissue> pairs with very low mutation counts (<=10) across binding site positions. **(b)** Number of mutations in accessible genomic regions for each tissue type (**Methods**). **(c)** Examples of mutation profiles for <TF-cluster, tissue> pairs at different q-value cutoffs. For each tissue type, the observed mutation counts are shown in color (using the same color scheme as in panels a and b), while the mean expected counts along with +/-1 standard deviation are shown in gray. Blue circles highlight the positions with q- value<0.1. Vertical dashed lines mark the regions defined as being bound by the TFs, i.e. +/- 10 base pairs around the motif center. **(d)** Heatmaps of the most significant enrichments for <TF, tissue> pairs, focusing on the TFs with the lowest versus the highest gene expression levels in each tissue type (**Methods**). Right: bar plot quantifying the percent of TFs, within each category, that show significant enrichment at a q-value cutoff of 0.1.

Our catalog revealed that mutation enrichment at TF binding sites is widespread. Each of the 15 tissue types in our study showed enrichment at one or more of the 276 TF clusters analyzed (**Fig. 2a**). For further analyses, we filtered out 1,369 <TF-cluster, tissue> pairs that had 10 or fewer mutations at all binding site positions (pairs shown in grey in **Fig. 2a**). Of the remaining 2,771 <TF-cluster, tissue> pairs, almost one third showed significant enrichment (q-value<0.1; blue dotted lines in **Fig. 2a**) within at least one binding site position (31.58%, or 875 pairs; **Fig. S3a**). At a more stringent cutoff (q-value<0.05), we found that 751 of the 2,771 <TF-cluster, tissue> pairs (i.e. 27.1%) showed significant enrichment. The number of enriched TF clusters varied widely across tissues, ranging between 219/276 (79%) for skin tumors and 2/118 (1.7%) for biliary tract tumors (**Fig. S3c,d**). As expected, we noticed a correlation between the number of TF clusters enriched within a tissue type and the total number of cancer mutations in accessible DNA regions in tumors from that tissue type (**Fig. 2a,b**; **Fig. S3b**). This suggests that, as additional mutation data becomes available, we can expect to identify even more TFs with significant enrichment within TF binding sites, especially for tumors with low mutation counts in our study.

Next, we compared the TF clusters that showed *elevation* of mutations within the binding site (**Fig. S1b**) versus the TF clusters that showed a significant *enrichment* of mutations within at least one binding site position (**Fig. S3c**). These sets did not overlap completely (**Fig. S3e**). In fact, more than 50% of the <TF-cluster, tissue> pairs that showed mutation elevation did not show a corresponding enrichment of mutations above background, indicating that the mere presence of elevated mutation counts within the binding sites of a TF does not imply that mutations are enriched compared to what we would expect based solely on the binding site sequences. Moreover, we found that for <TF-cluster, tissue> pairs that showed significant mutation enrichment but at only a single position within the binding site region, computing the overall mutation elevation over the entire binding site (relative to its flanks) can actually drown out the enrichment signal (**Fig. S3f**). Taken together, these results emphasize the importance of using our modeling framework to detect genuine enrichment of cancer mutations at TF binding sites, and thus identify TFs whose binding is likely to contribute to mutagenesis in regulatory DNA.

The results presented in **Figure 2a** are at the level of TF clusters, with a separate mutation profile for each <TF-cluster, tissue> pair. In order to assess the range of TFs showing mutation enrichment, and to prioritize TFs for further analyses, we assigned to each TF the mutation profile of the cluster it belonged to. For TFs present in multiple clusters (**Table S1**), we used the profile with the strongest enrichment (lowest q-value) within the binding site region. Among the resulting 9,825 <TF, tissue> pairs (655 TFs, 15 tissue types), 7,991 pairs had at least 10 mutations per position; we focused on these TFs for further analyses. In total, ∼35% of <TF, tissue> pairs (2,765/7,991) showed significant enrichment at q-value<0.1, and ∼30% of pairs (2,384/7,991) showed significant enrichment at q-value<0.05. This set of enriched <TF, tissue> pairs can be thought of as candidates for potentially mutagenic TF binding within that tissue type. The resulting TF-level heatmap is shown in **Fig. S4c**. As expected based on our TF cluster-level analyses (**Fig. S3c**), the number of TFs showing significant enrichment varied widely among tissue types (**Fig. S4a-b**), and it correlates well with the overall number of cancer mutations available for each tissue type analyzed here (**Fig. S4b)**. These results further underscore the widespread nature of somatic mutation enrichment at TF binding sites, and the potential to identify even more <TF, tissue> pairs of interest as more cancer mutation data becomes available.

### Transcription factors expressed at higher levels show stronger enrichment of somatic mutations at their binding sites

To gather further evidence that the significant enrichment of mutations observed for certain <TF, tissue> pairs is likely due to TF binding, we repeated our analyses for subsets of binding sites that are most likely to reflect true binding events in the cell. In particular, we focused on the TFs mostly highly expressed in each tissue of interest, and hypothesized that these TFs would show a larger fraction of enriched <TF, tissue> pairs compared to lowly expressed TFs. Our rationale is that highly expressed TFs will bind to a larger fractions of their putative genomic binding sites identified in open chromatin regions, compared to lowly expressed TFs, and thus any signals due to TF binding will be amplified.

In order to test this, we used gene expression data from the GTEx (The GTEx Consortium 2020) to group the 655 TFs in our study into highly-expressed and lowly-expressed subgroups (**Methods**). We then generated TF-level heatmaps separately for the high expression and low expression subgroups (**Fig. 2d**). Across all tissue types tested, more TFs in the high expression subgroup showed significant mutation enrichment (q-value<0.1) compared to TFs in the low expression subgroup (**Fig. 2d**, right). This overrepresentation of enriched TFs in the high expression subset was seen at different cutoffs of high versus low expression (**Fig. 2d, Fig. S4d**), consistent with our expectation that highly expressed TFs, which are more likely to be bound to a large fraction of their putative genomic binding sites, show stronger mutation enrichment.

### A pan-cancer, position-specific catalog of mutation enrichment at TF binding sites

The number and identity of enriched TF clusters varies across tissue types. The largest number was seen in skin tumors, the cohort with the largest number of mutations in our study (**Fig. 2a,b**), with 219 of the 276 TF clusters showing a significant enrichment (q-value<0.1; **Fig. S3c**). This is consistent with previous studies that showed strong mutation enrichment at TF binding sites in skin melanomas (Sabarinathan et al. 2016; Frigola et al. 2021; Wasserman et al. 2026). Both analytical and experimental evidence points to differences between TF families, as well as differences in the extent to which increased damage versus reduced repair contribute to the observed mutation enrichment at the binding sites of different TF families (Frigola et al. 2021; Sivapragasam et al. 2021; Duan et al. 2024). Our study provides further evidence for this enrichment and extends its characterization to individual nucleotides across a broad range of TF families, an important foundation for experimental validation given the base-specific nature of DNA damage and repair.

Our study also identified mutation enrichment across various non-melanoma tumors, which have not yet been analyzed at scale. Analysis of TF-induced mutagenesis in non-melanoma tumors has typically focused on certain TFs, with variation in how enrichment was defined (Katainen et al. 2015; Yang et al. 2018; Morova et al. 2020). By profiling a large number of TFs across tissue types with a unified approach, our catalog presents a more holistic view of the potential prevalence of this effect. Overall, we observe that mutation enrichment at TF binding sites is more widespread than previously appreciated. Beyond melanomas, we found that esophageal tumors, the second largest group by mutation count in our study, also showed high enrichment, with 133 TF clusters comprising 385 individual TFs showing statistical significance (q-value <0.1) (**Fig. S3c, Fig. S4a**). For 11 out of the 15 tissue types analyzed, enrichment was seen in >10% of TF clusters with sufficient mutations for our analysis, and enrichment was seen in >30% of TF clusters for the 5 largest cohorts (**Fig. S3,S4**). In the remaining 3 tissues, most TF clusters had too few mutations to detect enrichment (gray bars, **Fig. 2a**), but some clusters still showed enrichment. Further profiling, especially in these tumors with less mutation data available, will significantly improve our understanding of the range of this effect.

Since TF-DNA interactions and mutagenic processes act at the level of individual bases, modeling the impact of their interplay should ideally differentiate between different binding site positions. Previous investigations of TF-induced mutagenesis in melanomas indeed showed variation in this effect when assessed at different bases within a TF binding site (Poulos et al. 2016; Frigola et al. 2021). We reasoned that any assessment of this effect across cancer types must also be position-specific, and hence we performed our enrichment assessment at each binding site position separately. While such an approach has reduced power for some TFs, it nevertheless is more representative of the processes underlying an increase in the rate of mutagenesis at TF binding sites. By generating an observed count and an expected count at each binding site position, we observed that the mere presence of a high number of mutations at a position was not predictive of it being significantly enriched relative to what is expected by chance, based on the DNA sequence of the binding sites (**Fig. S3f**). Additionally, we found that in cancers with very few mutations, we still observe enrichment at some TFs (**Fig. 2a**). Since the total number of mutations in these cases is low, they often do not show a large increase within the entire binding site, despite having specific spots with high enrichment (**Fig. S3g).** Signals at these positions can be obscured by aggregating mutations over the entire binding site, especially if only 1 or 2 positions show large deviations. Moreover, for TFs that showed enrichment in more than one tissue type, the position (or positions) within the binding site where enrichment was seen can be different, (**Fig. 3a, Fig. S5**), indicating that they likely arose from different mechanisms. Taken together, our results highlight the importance of calculating deviations within the binding site in a position-specific manner, rather than by aggregating signals over the entire binding site.

**Figure 3:**
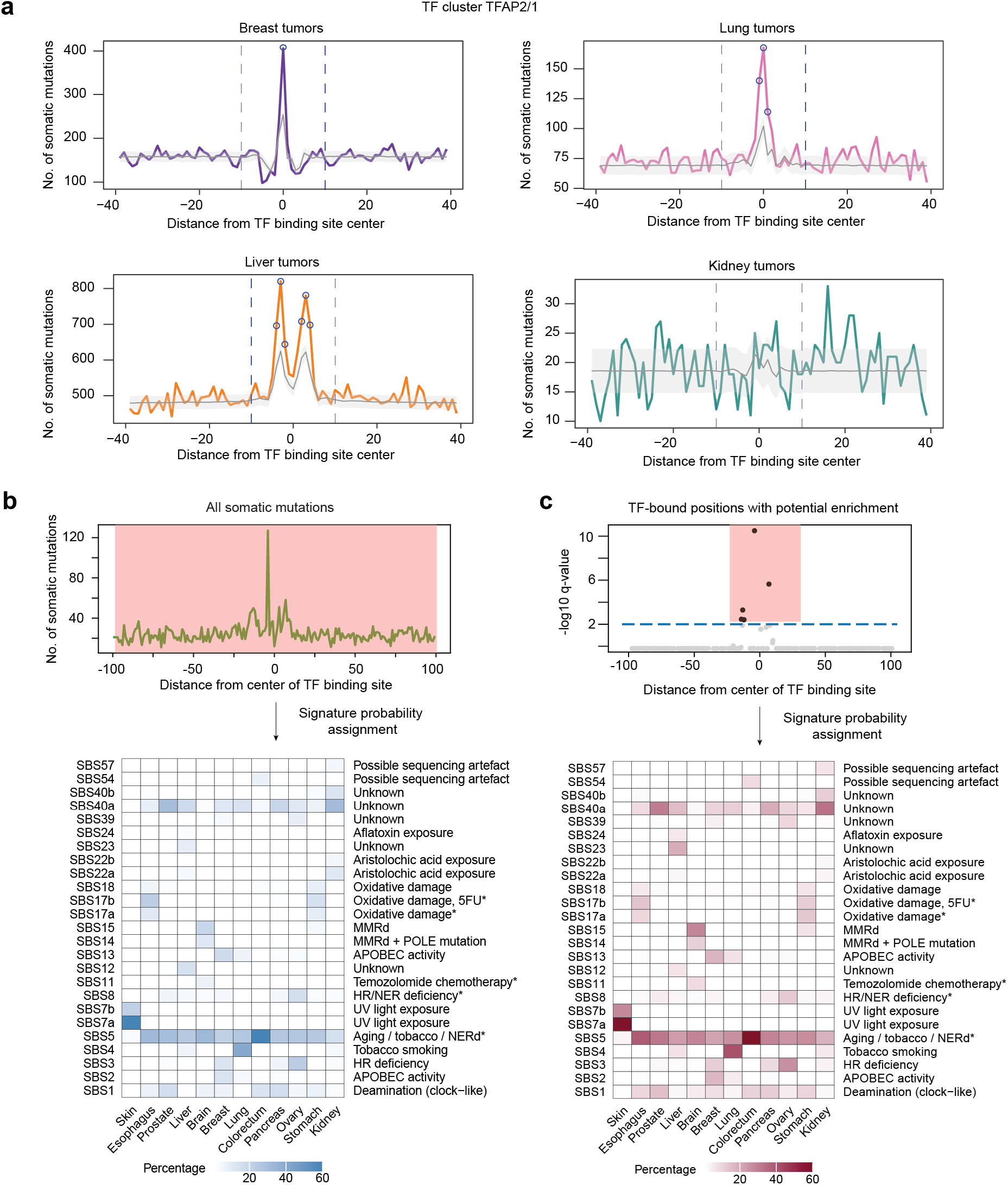
Mutational processes involved in potential TF-induced mutagenesis. **(a)** Mutation profiles (as described in Fig. 2) for the TF cluster TFAP2/1 across different tissues (breast, lung, liver, and kidney). Enriched positions (q-value < 0.1), highlighted in blue, differ across tissues. **(b)** Contributions of single-base substitution (SBS) mutational signatures to all somatic mutations in 12/15 tissues used in the study (**Methods**). Bone, uterus, and biliary tract tumors, which had fewer than 10 enriched TF-clusters, were not used for signature analysis. Each cell shows the percentage of a tissue’s mutations attributed to that signature. All signatures that contributed at least 5% of mutations in at least one tissue in (b) or (c) are shown. Etiologies are sourced from COSMIC, with asterisks (*) marking those with proposed mechanisms but unclear supporting evidence. **(c)** Same as (b), except that only mutations at enriched TF-binding site positions (q-value < 0.1) are included when calculating SBS contributions.

### Assessing potential mechanisms for TF-induced mutagenesis

TFs may show increased mutation rates within binding site positions if their interactions with DNA have downstream effects on mutagenic processes. This phenomenon of an observed increase in mutation rates due to TF binding, which we refer to here as TF-induced mutagenesis, can stem from increased DNA damage or decreased DNA repair within TF-bound DNA. Within our study, potential enrichment at binding sites was strongest in tumors from skin tissues, consistent with previous reports of TF-induced mutagenesis within melanomas. In melanomas, most mutations arise from damage due to ultraviolet (UV) radiation (Pleasance et al. 2010; Alexandrov et al. 2013), primarily repaired by the nucleotide excision repair (NER) pathway. Consequently, TFs have been implicated in mutagenesis within melanomas due to by (a) increase in UV damage due to TF binding (Elliott et al. 2018; Mao et al. 2018), (b) inhibition of NER of (Sabarinathan et al. 2016), or (c) by affecting both DNA damage and DNA repair processes (Duan et al. 2024; Wasserman et al. 2026). Beyond UV lesions, TF binding has been proposed to increase mutagenesis by reducing repair of lesions from endogenous cellular processes, such as abasic sites created during spontaneous depurination (Morova et al. 2020), and replicative DNA mismatches (Reijns et al. 2015; Zhu et al. 2025). Our pan-cancer catalog found significant increases in mutation rates for many TFs across tumors in a diverse set of tissue types (**Fig. 2a**), suggesting that additional DNA damage and repair processes might be implicated in TF-induced mutagenesis.

However, even when a specific hypothesis for TF-induced mutagenesis is known, experimental validation can be challenging. Such validation would ideally include perturbations to TF binding and to the specific DNA damage and repair processes implicated in the hypothesis. Isolating these interactions is further complicated by the binding of many paralogous TFs to similar DNA sequences, as well as the potential of multiple DNA damage and repair processes to generate a given mutation. Consequently, studies aimed at validating TF-induced mutagenesis mechanisms require carefully designed experimental systems, where confounding factors can be controlled. We recently designed such a system to demonstrate that TF binding can impede the repair of replication errors by the mismatch repair (MMR) pathway, resulting in increased mutation rates at TF binding sites (Zhu et al. 2025). Even with a single specific hypothesis, this required carefully chosen DNA sequences, TF proteins, and multiple knockout experiments to validate that binding increased mutation rates specifically through competition with MMR. To our knowledge, our prior study remains the only demonstration of TF-induced mutagenesis in a cellular system. Additionally, hypotheses of mutagenesis stemming from reduced DNA repair require TFs to bind to their binding sites even in the presence of DNA damages. We recently demonstrated, using novel high-throughput assays for protein-DNA binding, that TFs indeed bind to DNA with lesions, including mismatches resulting from DNA replication errors or deamination of methylated cytosines (Afek et al. 2020), and UV-induced DNA photoproducts (Mielko et al. 2023). However, our prior work also showed that TF binding to DNA lesions cannot be reliably predicted from experimental data and models of TF-binding to undamaged, Watson-Crick DNA (Afek et al. 2020; Mielko et al. 2023).

Taken together, these studies established that experimental validation of TF-induced mutagenesis requires novel experimental design to perturb the implicated DNA damage and repair pathways, as well as TF binding, while also requiring novel experimental assays to assess TF binding to damaged DNA substrates. These are not trivial tasks, and require significant time and resources. Consequently, assessing which TFs and DNA damage or repair processes ought to be prioritized for future studies is a major roadblock for experimental validation of TF-induced mutagenesis. Our position-specific pan cancer catalog represents a valuable resource to help identify and prioritize TFs and mutagenic processes for follow-up studies of mutagenesis induced by TF binding.

Interestingly, we noticed cases where the same TF-cluster shows mutation enrichment at different positions within the binding site, depending on the tissue type (**Fig. 3a**). This is consistent with previously observed patterns within hypermutated CTCF binding sites (Katainen et al. 2015; Poulos et al. 2016), and arises due to different damage and repair processes being active in tumors from different tissues. We reasoned that the enrichment seen across our catalog likely stems from distinct processes, and would require TF-specific annotation in order to be studied experimentally. To do this, we sought to assign putative mutagenic processes to the mutations used in our catalog, based on the mutational signatures present in the corresponding tumor samples (Alexandrov et al. 2013; Alexandrov et al. 2020).

Using the predefined set of single-base substitution (SBS) mutational signatures defined in COSMIC (Tate et al. 2019), we estimated their contributions to each tumor sample in our study. These activities, together with the SBS-specific mutation-type probabilities, were used to calculate the contribution of each SBS to each observed mutation in our samples. We then used the SBS-specific probabilities for each mutation to estimate the total number of mutations contributed by each signature within all tumor samples for each tissue type in our study (**Table S3**, **Methods**). These signature decompositions agreed well with previous studies, and identified high mutational loads for tissue-specific processes such as UV signatures (SBS 7a, 7b, 7c) in skin tumors, smoking-induced signatures (SBS 4) in lung tumors, etc. (**Fig. 3b**). Using these mutational loads, we then focused on mutations in positions identified as enriched in our catalog (q-value < 0.1). These specific positions showed similar distributions to the overall mutational burden for most tissues (**Fig. 3c**), suggesting that the processes involved in mutagenesis at TF binding sites are largely the same as those broadly active across the genome.

Assignment of signatures to the enriched mutations observed at TF binding sites can be used to ask which mutagenic processes are most active at the binding site positions which show mutation enrichment. Given the large number of TFs for which enrichment was observed in skin tumors, UV signatures were the largest source of mutations at these sites. This trend is consistent with prior work showing that both UV damage formation and repair of UV lesions is affected by TF-DNA binding (Frigola et al. 2021; Wasserman et al. 2026), making mutations more likely to hit binding site positions for TFs involved in these interactions. However, our pan-cancer enrichment map enabled the addition of several other potential sources of mutagenesis at TF binding sites, for which cancer- and mutagen-specific studies have not yet been conducted. Indeed, the majority of mutational signatures active genome-wide were also active at enriched mutation positions, suggesting that TF binding could interact with other common DNA damage and repair processes, by modulating damage formation and/or repair.

Broadly, the non-UV signatures enriched at TF binding sites can be grouped into two classes: a pan-cancer class that is active in most tissue types (**Fig. 4a**), and a tissue-specific class that is active in only one or a few tissues (**Fig. 4b**). The first class contains signatures such as SBS1, SBS5, and SBS40, which typically correlate with age. Together, these mutations contributed 52.5% (41-84% per tissue type) of enriched position mutations in non-skin tissues. The mechanisms behind SBS5 and SBS40 are currently unknown, but their clock-like and pan-cancer nature suggests that these are due to common cellular processes. SBS1 is known to come from one such mechanism, through the spontaneous deamination of methylated cytosines (Duncan and Miller 1980; Pfeifer 2006; Nik-Zainal et al. 2012). This deamination yields thymine, leading to a G-T mismatch that is typically repaired by the base excision repair pathway (Walsh and Xu 2006; Jacobs and Schar 2012). Recently, a role for the mismatch repair pathway in correcting this mismatch has also been proposed (Poulos et al. 2017). Inhibition of these repair processes by TF binding could lead to a higher proportion of uncorrected G-T mismatches, which in turn would lead to increased rates of C>T mutations.

**Figure 4:**
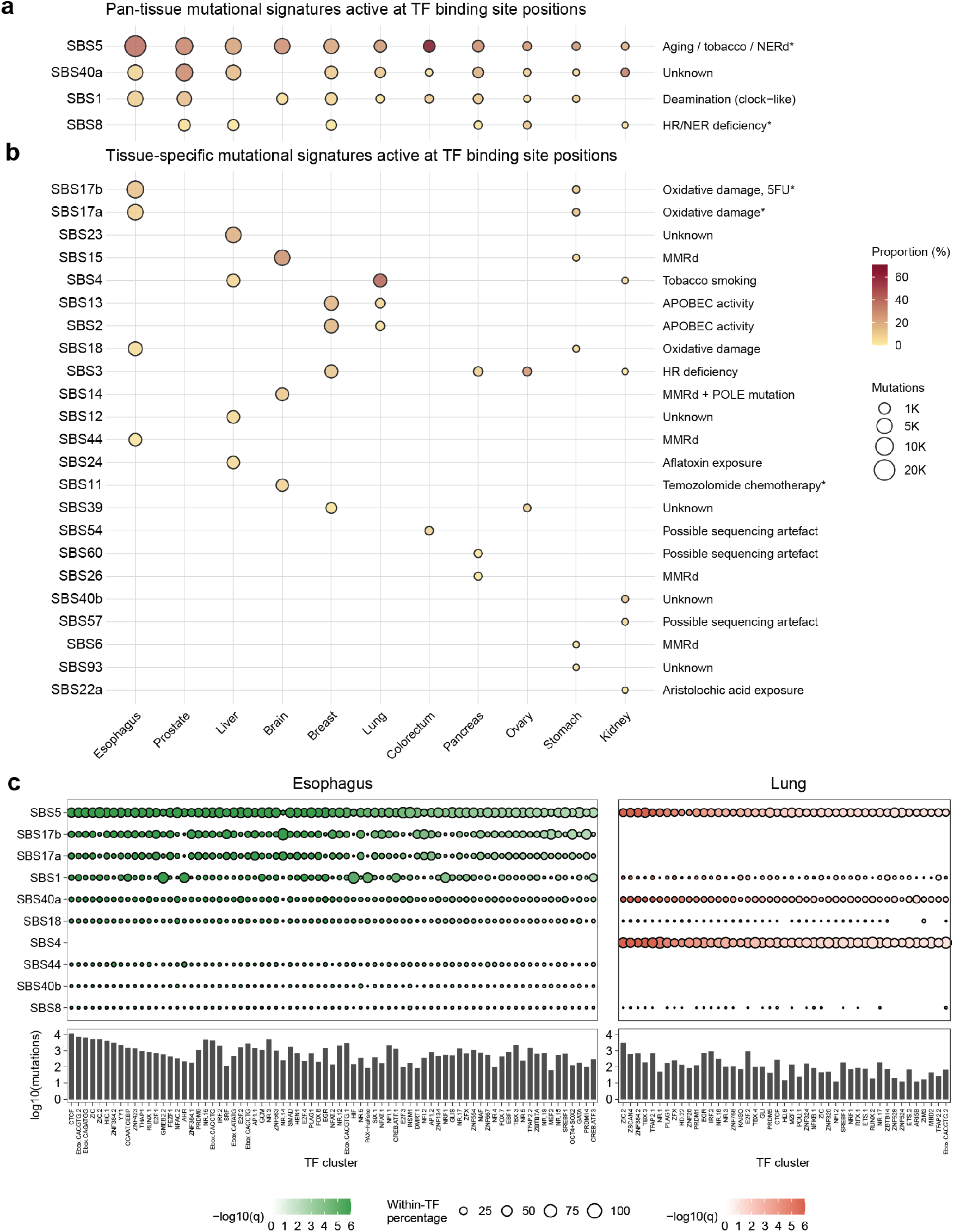
Attributing enriched mutations to mutagenic mechanisms. (a,b) Signature contributions at enriched TF binding sites positions in non-skin tissues, shown as number of enriched- position mutations attributed to each SBS signature (circle size) and percentage of each tissue’s enriched-position mutations it accounts for (circle color), split into (a) pan-cancer signatures and (b) tissue-specific signatures. Pan-tissue signatures were present in at least 6 of the 11 tissues shown, with a signature counting as present if it contributed to at least 3% of enriched TF binding site mutations. **(c)** Signature composition at individual TF clusters for esophageal tumors (left) and lung tumors (right). TF clusters with at least one enriched binding-site position are shown, ordered by strength of enrichment (q-value). For esophageal tumors, the top 75 TF clusters are shown. The top 10 signatures contributing to enriched TF binding site positions in lung and esophagus are shown.

Some mutational signatures showed high activity only in specific tissue types (**Fig. 4b**). For example, signatures proposed to be associated with oxidative damage, such as SBS17a, SBS17b, and SBS 18, contributed 12% of mutations at enriched TF binding site positions, largely due to high activities at these positions in esophageal tumors. Other tumor-specific mechanisms included the tobacco smoking signature SBS4, which was a large contributor to mutagenesis at enriched sites in lung cancers, but not in tumors from other tissue types. Similarly to UV damage, these patterns could result from an increase in oxidative or tobacco damage at TF binding sites, or from a reduction in the repair of the corresponding lesions (if the lesions are bound by TFs strongly enough to compete with the repair proteins that must recognize these lesions and initiate their repair).

Overall, signature decomposition can provide clues regarding the mutational mechanisms that might be associated with mutation enrichment at TF binding sites. The diverse nature of these mechanisms suggests that TF binding can cause an enrichment of mutations by interfering with diverse damage sources and repair pathways. Similar to our pan-cancer map of mutation enrichment, we also provide a pan-cancer map of mutational signatures contributing to the enriched mutations, depicted for each cancer type separately (**Fig. S6-S16; Table S4**), with esophageal and lung tumors shown in **Fig. 4c**. The differences in active signatures across tissues again underline the importance of our tissue-specific approach to analyzing mutation enrichment at TF binding sites.

### Connecting mutation enrichment profiles to experimental pathways for validation

By building a pan-cancer map of mutation enrichment at TF binding sites, our goal was twofold: 1) to provide insights into the magnitude and diversity of TF effects on mutagenesis, and 2) to provide testable hypotheses about TF-induced mutagenesis that can be validated through mechanistic studies. Our catalogs of enrichment profiles and mutational signatures in TF binding sites fulfill the first goal. Below, we discuss ways to derive testable hypotheses for increased mutagenesis at transcription factor binding sites using this roadmap.

In **Fig. 5a** we show the mutation profile in esophageal tumors for the TF cluster AHR, with a binding site position that shows significant enrichment of mutations. At this position, we saw a large number of mutations attributed to SBS1, the deamination signature discussed above. Even though this signatures was active in the flanking regions as well, those positions did not show similar enrichment. Combined with the signature etiology, and our recent work showing that TFs can bind strongly to mismatched DNA (Afek et al. 2020), our analysis allows us to derive specific hypotheses to understand hypermutation connected to TF binding (**Fig. 5b**), namely, in this case, that G-T mismatches bound by AHR could go unrepaired by base excision repair (BER) or mismatch repair (MMR) pathways, by preventing recognition of the G-T mismatch lesions by BER or MMR repair proteins (such as MBD4, TDG, or MutSα).

**Figure 5:**
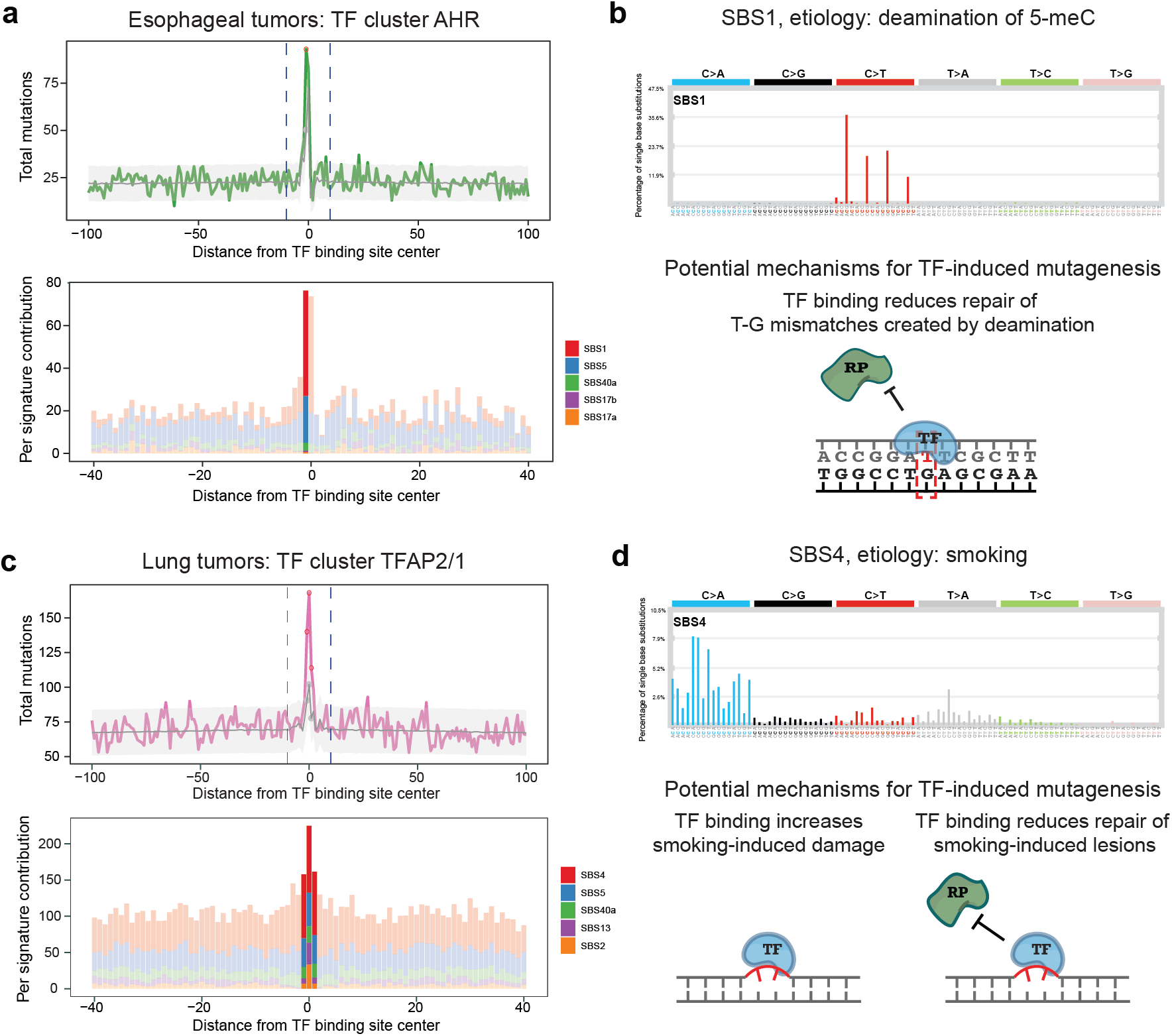
Putative mechanisms for increased mutagenesis at TF binding sites. **(a)** Mutation enrichment at the AHR TF cluster in esophageal tumors, along with SBS contributions. Highlighted positions within signature plots correspond to enriched positions, and the gray line represents the expected mutation count. Contributions are shown for the top five SBS signatures with the highest contributions in the profiled window (+/- 40bp around the motif center). **(b)** Etiology for the top signature, SBS1, present at AHR sites and the corresponding mechanisms through which the observed enrichment may occur. **(c)** Same as panel a, for the TFAP2/1 TF cluster in lung tumors. **(d)** Same as panel b, for SBS4, the signature with the highest contribution at the TFAP2/1 binding site positions with significant mutation enrichment. Accessibility sets for binding sites for these figures were: ATAC-seq for AHR and DNase-seq for TFAP2/1.

In lung tumors, we found large contributions from SBS4 (**Fig. 4c**). At specific TF binding sites, this can be visualized by highlighting enriched positions within TF binding sites for which high contributions come from SBS4 (**Fig. 5c**). Similar to the example above, SBS4 is present elsewhere in the binding site, but does not always result in enrichment, highlighting the base-specific nature of hypermutation patterns. SBS4 is attributed to tobacco smoking (Nik-Zainal et al. 2015), creating predominantly C>A mutations. Tobacco smoke contains DNA-damaging agents such as benzopyrene (BaP), which can form adducts on the N2 position of guanines. Such adducts are typically removed by the nucleotide excision repair (NER) pathway. Inhibition of this pathway by TF binding leads to widespread enrichment of UV-induced mutations in melanomas, and it might be responsible for mutation enrichment in lung tumors as well. Our analysis identified specific TFs (such as AP2 factors, **Fig. 5c**), and TF binding site positions that should be prioritized for future mechanistic studies of the interplay between human TFs and smoking-induced damage and repair (**Fig. 5d**).

Thus, our position-specific analysis of TF binding site mutations and their concurrent signature contributions can help uncover sources of TF-induced mutagenesis. We provide signature decompositions for all enriched <TF-cluster, tissue> pairs (**Table S4**), which will assist in nominating further hypotheses that can be tested experimentally.

## DISCUSSION

Mutation rates show high variation across the genome, reflecting underlying differences in the activities of DNA damage and repair processes. Recent work has suggested that TF binding sites in regulatory DNA can show localized hypermutation patterns, but the overall prevalence and mechanistic understanding of this effect remain unclear. In this work, we generated a comprehensive pan-cancer catalog of mutation rates at TF binding sites for tumors from 15 tissue types and compared the observed mutation rate within these binding sites to the expected mutational burden. This was done using a rigorous statistical approach that analyzes each TF binding site position separately, and accounts for sequence-dependent and local (background) variation in mutation rates. By comparing the observed mutation counts to the expected background mutation counts, we were able to identify whether specific positions within TF-bound sites showed a statistical enrichment for somatic mutations. We found that mutation enrichment at TF binding sites is more widespread than previously reported, and extends to tissue types not previously analyzed. These patterns are highly TF- and tissue-specific, showing that analysis of active mutational processes within regulatory DNA should be performed for each TF and tissue type independently. Consistent with tissue-specific differences in mutagenic mechanisms at TF binding sites, we observed that positions within the TF binding sites that show enrichment can differ across tissues. We also observed that a high mutation burden alone is not necessarily associated with overall enrichment, which underlines the importance of a statistical approach that considers each binding site position separately.

All 15 tissues analyzed in this study showed mutational enrichment at TF binding sites, albeit with widespread heterogeneity. As more somatic mutation data becomes available, the nature of TF-induced mutagenesis can be better characterized for tissues with low mutation counts in our study. Recently, co-binding of associated TFs was also reported to modulate the nature of mutational profiles observed at TF binding sites in liver cancer, along with differences between enhancers and promoters, indicating that local chromatin also shapes the nature of TF-induced mutagenesis (George et al. 2026). Further incorporation of such local differences, while comparing mutational patterns at TF binding sites, are also warranted as more mutation data and chromatin profiling data are generated.

Previously, TF-induced mutagenesis has been well characterized within melanomas, where TF binding is thought to: 1) compete with nucleotide excision repair, and 2) increase the amount of UV damage at the binding sites of certain TFs (Sabarinathan et al. 2016 ; Frigola et al. 2021; Wasserman et al. 2026). However, our results show that such patterns exist across various other tissue types, and associate with diverse mutational signatures, suggesting that other interactions between TF binding, DNA damage, and DNA repair can also produce hypermutation patterns for specific TFs. Signatures associated with other damage sources such as tobacco adducts and oxidative damage, and those associated with repair of deamination-induced mismatches, are likely to contribute towards the mutation patterns seen at TF binding site positions with significant enrichment. By annotating enriched TF binding site positions, together with the active mutagenic processes that are likely acting at those positions, we provide a systematic approach for prioritizing these interactions for further mechanistic studies. We believe that our catalog can enable the identification of TFs whose DNA binding can incur a mutagenic cost, based on their interactions with specific damage or repair processes.

We recently showed experimental evidence for such a TF-induced mutagenesis mechanism, whereby multiple TFs were found to impair the recognition of DNA mismatched lesions by MMR enzyme MutSα, resulting in a higher mutation burden at TF binding sites (Zhu et al. 2025). This required an experimental design which depended on the selection of specific TFs, DNA lesions, and repair processes (in this case, MMR). Our current work expands the number of TFs and mutagenic processes for which similar tests can be undertaken experimentally. However, this requires further conceptual and technical advances in the profiling of DNA damage and repair processes across the genome. Several techniques enable the mapping of both UV lesion formation and NER activity using next-generation sequencing (Hu et al. 2015; Mao et al. 2016), and have consequently been used to study TF-induced mutagenesis in melanomas at higher resolution. Apart from UV lesions, recent techniques have also been designed to profile other lesion types and repair intermediates (Ding et al. 2017; Poetsch et al. 2018; Tang et al. 2022), though these techniques differ in their conclusions and often lack the single base-pair level resolution necessary for analyses within TF binding sites (Pfeifer and Jin 2024; Wasserman et al. 2026). Further refinement of techniques for high-resolution mapping of DNA damage and repair are needed in order to validate TF-induced mutagenesis in cellular systems.

Another aspect that requires further characterization is the binding of TFs to damaged DNA. For TFs to impede the repair of DNA damages, they must first outcompete the repair enzymes that recognize the DNA lesions to initiate repair. We recently profiled the effects of UV-damage on TF binding (Mielko et al. 2023) and showed that this competition requires first assessing the effect of damage on the binding of each specific TF. We also recently showed the widespread effects of mismatches, which are a major source of somatic mutations (Tomasetti et al. 2017), on TF-DNA recognition (Afek et al. 2020). Crucially, the effect of mismatches was not explained by knowledge of Watson-Crick binding for the TFs tested. Together, these studies show that assessing hypermutation due to TF binding require experimental designs targeting specific TFs and repair pathways. Our study can power future analyses in this direction by identifying viable candidates for TF-repair enzyme competition.

The mutagenic cost of TF binding has important consequences for the study of cancer genomes. A key challenge in the field is to identify non-coding drivers in regulatory DNA, where signals of selection are weak. The largest pan-cancer study identifying non-coding drivers combined several techniques for driver discovery (Rheinbay et al. 2020), but current methods are not equipped to capture localized sources of hypermutation (Elliott and Larsson 2021). This hypermutation can increase the rate of certain passenger regulatory mutations, which might then show recurrence (Kaiser et al. 2016; Buisson et al. 2019). As a result, the Rheinbay et. al study manually flagged driver candidates that might result from local hypermutation – defined as UV mutations due to TF binding in melanoma, AID activity in lymphomas (Robbiani et al. 2009; Puente et al. 2015), and APOBEC mutagenesis in several cancers (Buisson et al. 2019). Further identification of driver mutations would require the profiling of an even larger set of cancer genomes, and in a fundamentally low signal-to-noise paradigm, such annotation would continue to be vital. Here, we present evidence that TF binding has a more widespread mutagenic role than the previously described role in melanomas. This finding can help identify similarly recurrent mutations in binding sites that are passengers, or aid statistical approaches that seek to account for localized sources of variation (Guo et al. 2020). The heterogeneity we see within TFs for most non-skin tumors, as well as heterogeneity across tumors for the same TF (**Fig. 2a,d**), suggest that these approaches should model the binding sites of each TF and each tumor type separately.

Additionally, our results have implications for studies of germline variation and natural selection. In our study, we focused on somatic mutations and we identified processes through which TF binding could lead to mutagenesis in cancer cells. However, TF-induced mutagenesis could also operate in the germline and confound existing models of natural selection in the regulatory genome, which assume similar rates of background mutation within TF binding sites and flanking regions (Torgerson et al. 2009; Arbiza et al. 2013). Recent studies provide support for TF-bound mutagenesis operating in the germline (Dukler et al. 2022; Seplyarskiy et al. 2023), but further characterization would require TF-specific tests similar to those undertaken here for somatic mutations in tumor genomes.

## ACKNOWLEDGEMENTS

This work was supported by National Science Foundation (NSF) grant #2529591 (to R.G.), and National Institute of General Medical Sciences (NIGMS) grant R01-GM135658 (to R.G.). ChatGPT (OpenAI) was used for minor language edits to correct typographical errors and improve clarity and flow. All edits were reviewed by the authors, who take full responsibility for the final text.

## METHODS

### Cancer somatic mutation data

The mutation data used in this study was obtained from the International Cancer Genome Consortium (ICGC), release 28. Files containing simple somatic mutations for the 36 projects listed in **Table S1** were downloaded from the ICGC data portal (Zhang et al. 2019) and filtered for single base substitutions identified in whole genome sequencing data. Next, we grouped the ICGC projects by tissue to increase the number of somatic mutations available for our analyses, as described in **Table S1**. All subsequent analyses were done at the level of individual tissues. As more data becomes available from studies that include larger numbers of donors, our analyses can also be applied in a more fine-grained manner, focusing on specific tumor types or subtypes. As shown in our manuscript, the power to identify TF binding site positions with a significant enrichment in somatic mutations is generally proportional to the size of the mutation data sets used, as expected.

### Transcription factor binding site identification

Binding sites for each tissue type were identified using open chromatin data from: 1) tissues or relevant cell lines (DNase-seq: Roadmap Epigenomics Project (Roadmap Epigenomics et al. 2015) and ENCODE (Thurman et al. 2012)), and 2) tumor samples (TCGA (Corces et al. 2018)). The source datasets for each tissue/tumor are listed in **Table S1**. Here, we chose to use both data types (ATAC-seq in the tumors and DNase-seq in the healthy tissues) because it is not entirely clear whether focusing on accessible regions in the healthy tissues versus the tumors is most relevant for studies of mutagenesis in regulatory DNA. Chromatin accessibility changes during tumorigenesis. At the same time, for any individual tumor analyzed for somatic mutations, it is unknown when each mutation occurred during the progression from healthy cells to tumor cells. For this reason, when performing our analyses to identify significant enrichments of mutations in TF binding sites, we call sites either in: 1) DNase-seq peaks in healthy cells; 2) ATAC-seq peaks in tumor cells; 3) peaks that overlap between the two types of data; and 4) the union of all peaks from the two types of data. We consider a position within TF binding sites to be significant based on its minimal p-value across the four analyses.

For each tissue type, whenever multiple DNase or ATAC seq peaks sets were available (i.e. multiple cell lines or Roadmap projects, or multiple tumors in TCGA), we used the merge functionality from the BEDtools suite (Quinlan and Hall 2010) to obtain one DNase-seq and one ATAC-seq accessibility file per tumor/tissue. For consistency between the two types of accessibility data, the 501-bp, hg38 ATAC-seq peaks reported in (Corces et al. 2018) were trimmed to be 150 bases (similar to the typical range for the DNase-seq datasets used) and converted to hg19 using the liftOver tool from UCSC.

Putative binding regions were identified using motif clusters from (Vierstra et al. 2020), for 276 of the reported 286 clusters. Some clusters had identical labels (e.g. SMAD), in which case we used the cluster with the larger number of motif matches reported. KLF and GC-tract clusters were not used due to the very large number of motif matches reported (as many as ∼50 million). Two clusters (HD/24 and ZNF431) were removed because they contained only mouse motifs with no corresponding human orthologous TF. For each cluster, genome-wide motif matches to the archetype cluster motif (Vierstra et al. 2020) were intersected with the open chromatin datasets described above. Each TF cluster corresponds to multiple individual TFs with similar motif preferences, typically members of the same structural family. The analyses reported in this study were done at the level of TF motif clusters, to ensure enough mutation counts for statistical tests within each cluster. As binding data for individual TFs becomes available in the relevant tumor/tissue types analyzed here, our analyses can be refined to focus on TF-specific profiles.

### Mutation profiles

The mutation profiles shown in **Fig. 1a,b** and **Fig. S1** represent aggregate mutation rates across all TF motif clusters. To generate these profiles, binding sites for individual TF motif clusters were merged and then intersected with accessible DNA regions (according to DNase-seq and/or ATAC-seq data) to create a unified single set of putative TF binding sites for each tissue type. The center of each merged site was labelled as the TF binding site center, then mutations were counted at each position for 400 base pairs on each side and divided by 1000 to obtain mutations per kilobase. The elevation of mutation rates in binding sites (**Fig. S1a**) was calculated by computing mutation counts within the binding site (+/-10 base pairs from the center) and the flanking regions (150-300 base pairs from the center). For each tissue type, the elevation numbers depicted in **Fig. S1a** were calculated by finding the relative increase in the number of mutations seen at the binding site, compared to that seen in the flanks. To identify TF motif clusters that showed elevation, we then calculated the number of individual clusters that had a binding site mutation rate increase of more than 10%, compared to the flanking regions. TF motif clusters that had fewer than 10 mutations in either the flanks or the binding site were removed from this analysis. We then repeated this analysis at the level of individual TFs, by mapping each cluster to its constituent TFs (**Table S1**). For TFs that belonged to multiple TF clusters, the highest elevation seen across the clusters was reported.

To build mutation profiles for each TF, we first collected somatic mutation data for the ICGC projects listed in **Table S1**. We built a separate mutation profile for each TF cluster in a given tissue by using the binding sites called in that tissue (using the archetype cluster motif, as described above), intersected with the corresponding cancer somatic mutation data sets. Once a mutation profile was built for a particular TF motif cluster, that profile was used for all paralogous TFs mapped to that cluster (**Table S1**).

For each <TF-cluster, tissue> pair, mutation profiles were built by computing a position-specific mutation rate for each nucleotide within the binding site and flanking regions, defined as 100 bases from the center of the binding site on either side. For each base in this window, we computed the total number of times a mutation was observed at that spot, by aggregating the counts across all binding sites. For these analyses, the center of the binding site was assigned position 0. We then calculated a background rate for each position within the window of interest, as described below.

### Background mutation rates

For each nucleotide position in a mutation profile, we computed a background rate based on the trimer content at that position. First, we calculated the mutation rate for each of the 64 possible trimers, per tissue type. For each trimer, we counted the number of times mutations were observed in that trimer context, with the mutation occurring at the central base. This yielded an expected mutation rate for each trimer, defined as:

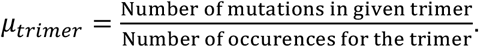

We calculated this rate by considering both trimer and mutation counts only in open chromatin regions, where the vast majority of active TF binding sites are located. These rates were computed separately for each tissue, using tissue-specific open chromatin regions and somatic mutations, and the mutation rates for a trimer and its reverse complement were combined by taking their average. Next, we modeled the occurrence of a mutation at a specific observed trimer by a Bernoulli process, where the probability of the mutation is given by the expected trimer mutation rate. Under this assumption, we can write the probability that an observed trimer is mutated as: *p_trimer_* = *μ_trimer_*. For each collection of binding sites, we built background mutation rates as described below.

Using the expected trimer rates, we calculated the number of expected mutations at each position in the window of interest (+/- 100 base pairs from the binding site center). To do this, for each position *i* we identified the observed trimer sequences (centered on position *i*) in each of the *n* binding sites under consideration. For position *i*, we then have *n* Bernoulli trials, with different probabilities of observing a mutation. These can be written in terms of the expected trimer rates as: *p_i_*_,j_ = *μ_i_*_,j_, where *j* = 1,2,3, . . . *n*, and *μ_i_*_,j_ represents the trimer at position *i* for binding site *j*.

Under the null/background model, there is no effect of TF binding on the expected rates in TF binding sites. Thus, we can assume that mutations occurring at different binding sites are independent of each other, and consider that each of the *n* Bernoulli trials described above are independent. We can then model the total number of expected mutations at position *i* as a Poisson-binomial process, which is the probability distribution of a sum of independent Bernoulli trials. For this distribution, the mean (and the variance) are given by the sum of the means (and variances) of the *n* Bernoulli distributions. To visualize the expected mutation counts, we computed a mean and a variance for each position, under the Poisson binomial assumption. This is given by:

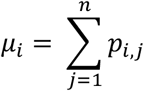

for the mean, and

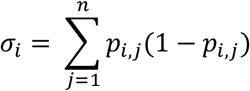

for the variance, following standard properties for a Bernoulli process.

We then visualize the mutation profiles by calculating the observed number of mutations and the mean and variance (+/- 1 std. deviation in the plots) for the expected/background mutation counts, for each of the 201 nucleotides in the windows being considered. This yields, for each TF motif cluster (and thus each TF represented in the motif cluster), a mutation profile in the tissue being profiled.

Our procedure to generate expected mutation counts at TF binding sites and their immediate flanking regions also accounts for local variation in mutation rates across the genome. Since the expected trimer rates are computed from all accessible genomic sequences, they are not tailored specifically to the local context of each TF of interest. For example, the binding sites of some TFs may be enriched in promoter regions of highly expressed genes, or in early-replicating regions, or in regions with a particular chromatin environment. Such broader genomic features can affect the local mutation rates at and around TF binding sites, independently of direct TF binding. In order to correct for variation seen among different regions at the local level (due to cellular processes such as replication, transcription, chromatin environment, etc.), previous work on mutation rate variation around TF binding sites typically normalizes the expected mutation rate to the local window around the TF being examined (Katainen et al. 2015; Vierstra et al. 2020). In other studies, which rely on simulations, the total number of mutations simulated in the window of interest is set to the observed mutation count in that window, in order to yield expected mutation rates (Poulos et al. 2016; Sabarinathan et al. 2016). We followed a similar approach to account for local differences in mutation rates, which can affect studies of hypermutation at TF binding sites. In our calculations, for each TF (or TF motif cluster) we first computed expected mutation counts using the procedure described above, namely by summing up expected mutations in the trimers observed at the TF binding sites and their flanking regions (**Fig. S2a**). These expected counts were generated for each position in the 201-nucleotide window (100bp on each side of the TF binding site center). Next, the flanking regions of the binding sites, defined as 30bp to 100bp from the binding site center on each side, were used to compute an average expected mutation count in the flanks, which was then scaled, by a constant scaling factor, to match the average observed mutation count in the same flanking windows. This scaling factor was applied to the mutation probability of each trimer, and then the expected mutation counts were recomputed, using the adjusted trimer mutation rates, across the entire 201-nucleotide region including the TF binding site (**Fig. S2b**). This procedure allows us to assess the potential effects of TF binding specifically, since only the flanking regions right outside the TF binding sites are used for normalization. After normalization, average expected counts and average observed counts are the same in the flanks, as expected (**Fig. S2c**, left plots), but they might differ in the binding site region (**Fig. S2c**, right plots), depending on the precise TF and tissue type considered. Next, we identify positions within TF binding sites where the observed mutation count is significantly higher than the expected count, as described below. At these positions, given that sequence-induced variation and local variation in mutation rates are accounted for, we hypothesize that TF binding is likely to contribute to the increased mutagenesis.

### Calculating position-specific mutation enrichment

To identify candidate positions that are enriched for mutations due to TF binding, our null hypothesis is that at a given nucleotide position, TF binding has no effect on the observed mutation count, i.e., that the observed count is simply the result of drawing a sample from the background mutation process, which we modeled as a Poisson binomial as described above. Under this assumption, the p-value for observing *k* mutations at a given position *i* can be written using the Cumulative Distribution Function (CDF) of the Poisson-binomial process at that position:

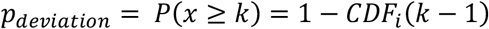

While the means and variances of the Poisson binomial distribution are simple to compute, as described above, exact CDFs are difficult to derive. To estimate these CDFs, we used the exact and approximate forms derived and implemented in the R-package poibin (Hong 2013). The package performs both exact and approximate calculations for Poisson-binomial CDFs, using a Refined Normal Approximation (RNA) (Volkova 1996) for approximate calculations, and a Discrete Fourier Transform (DFT-CF) approach for exact calculations. For efficient computation, we used the RNA approximation, which is expected to be precise for a large number of trials. While this approximation is fairly robust, we noticed differences in p-values computed using the exact and approximate versions for very small values of p (**Fig. S2e**). Thus, for p-values that were smaller than 10^-4^, we performed exact inference using the DFT-CF method.

Thus, for each position in each of 201-bp window analyzed, we compute: a) an observed mutation count, b) an expected mutation count, and c) a p-value to judge if the observed count is significantly higher than the expected mutation count. To account for multiple hypothesis testing, we use the Benjamini-Hochberg correction (Benjamini and Hochberg 1995) to correct for multiple hypotheses (with each position having a corresponding null hypothesis test) within the tested window (+/-100 bp from the center of the TF binding sites identified for a particular <TF-cluster, tissue> pair).

### Identifying TFs with significant enrichment in somatic mutations at their binding sites

For each <TF-cluster, tissue> pair, we then focused on the TF binding site region, defined as +/- 10 bp from the center of the 201-bp window, and we searched for the position with the minimum q-value, i.e. FDR-corrected p-value. All TFs (or TF clusters) with a minimum q-value < 0.1 in this analysis were considered to have a *significant* enrichment of mutations within their binding sites.

All putative TF binding sites used in our analyses were selected from accessible DNA regions of the human genome. Since most tissue types had multiple accessibility datasets available, we performed the analysis described above at various levels of stringency for DNA accessibility calls.

For each <TF-cluster, tissue> pair for which multiple accessibility datasets were available, we identified four sets of putative TF binding sites using: 1) only ATAC-seq peaks from tumor samples, 2) only DNase-seq peaks from tissue samples or relevant cell lines, 3) peaks found in both ATAC-seq and DNase-seq datasets for that tissue (the most stringent criterion), and 4) peaks found in either ATAC-seq or DNase-seq datasets for that tissue (the most permissive criterion). We performed the analyses in this manner because there are known differences between tumor and tissue accessibility that can affect somatic mutation rates (Polak et al. 2015; Corces et al. 2018). Since we cannot know, for the somatic mutations of interest, when they occurred along the tumorigenesis timeline, it is unclear whether the accessibility in the tissue of origin or the accessibility in the tumor cells is the most relevant for our analyses. Thus, to assess the potential effects of TF binding on mutagenesis broadly, we performed the analyses using the four accessibility modes described above (**Data Sets 1-15**). For further analyses and visualization (**Fig. 2**), we selected for each <TF-cluster, tissue> pair the accessibility mode for which the binding site region showed the largest deviation from the expected mutation profile, i.e. the smallest q-value (**Table S2**). This was done for each <TF-cluster, tissue> pair, as described, using all 276 TF motif clusters and 15 tissue types.

### Analysis of highly-expressed TFs

To analyze mutation profiles at the level on individual TFs, rather than TF clusters, we generated TF-level heatmaps (**Fig. S4**) by considering the mutational profile for each TF to be that of its TF motif cluster. For TFs that belonged to multiple clusters, the cluster with the most significant enrichment (lowest q-value) was chosen. This resolved the heatmap of <TF-cluster, tissue> pairs in **Fig. 2a** into a heatmap of <TF, tissue> pairs, which was used for the expression analyses. To identify highly expressed TFs in each tissue, we used RNA-seq data for adult tissues from the GTEx portal (Consortium 2020). For 14 of the 15 tissue types, GTEx v10 provided gene-level transcripts per million (TPMs) per tissue (median values across all donors). GTEx data was not available for bone, so we used RNA-seq data from the ICGC portal for the BOCA-FR project. This data was similarly converted into gene-level TPMs (mean across donors). This yielded, for all 15 tissue types of interest, a per-gene TPM value that was then used to determine gene-expression levels for the 655 TFs included in our study. For each tissue type, the 655 TFs were split into a top 25 percentile subset (High expression) and a bottom 25 percentile subset (Low expression). We then calculated, for both subsets, the percentage of TFs that showed enrichment (i.e., min. binding site q-value < 0.1). The tissue-wise comparison of this metric is depicted in **Fig. 2d**. The same analysis was repeated with top 10 percentile and bottom 10 percentiles, and is depicted in **Fig. S4d**.

### Mutational signature attribution

Single-base substitution (SBS) signature activities were calculated using SigProfilerAssignment from SigProfilerSuite (Diaz-Gay et al. 2023), using the COSMIC v3.4 reference set of SBS signatures (Tate et al. 2019; Alexandrov et al. 2020), applied to mutations within each included ICGC project. Projects were then mapped to tissues using **Table S1**. Next, for each individual somatic mutation we calculated a probability distribution over all SBS signatures, by multiplying the activity of a given signature with the probability of observing a mutation of the same type (trinucleotide context) under that signature. SBS-signature contributions were then calculated by adding up all the fractional probabilities for a given project or tissue. Tissue types with a small number of TFs enriched (bone, biliary tract, and uterus) were excluded from this analysis.

Overall SBS contributions were calculated as percentages for each per tissue type, for: (a) all somatic mutations included in the study, and (b) mutations falling within enriched TF-bound positions as defined above (**Fig. 3b,c, Table S3**). For further analysis of SBS signatures at enriched TF binding site positions, pan-tissue signatures were defined as signatures detected at more than 3% contributions across a majority of the included tissues, while the remainder were called ‘tissue-specific’ signatures (**Fig. 4a,b**). For each tissue type, we then analyzed TF clusters with at least one significantly enriched binding site position (q-value < 0.1), and summed up signature contributions for the observed mutations within those positions (**Fig. 4c, Fig. S6-16, Table S4**).

## CODE AVAILABILITY

Code used in the manuscript is available at https://github.com/harshitsahay/TFBS_Mutations

