## Supplemental Materials for "A pan-cancer catalog of mutagenesis at transcription factor binding sites"

### Supplementary Information for:
### A pan-cancer catalog of mutagenesis at transcription factor binding sites

Harshit Sahay[1,2,3,4], Yuncheng Duan[5], Raluca Gordân [2,3,5*]

[1] Program in Computational Biology & Bioinformatics, Durham, NC 27708, USA
[2] Center for Advanced Genomic Technologies, Durham, NC 27708, USA
[3] Department of Biostatistics & Bioinformatics, Durham, NC 27708, USA
[4] Computational and Systems Biology Program, Sloan Kettering Institute, Memorial Sloan Kettering Cancer Center, New York, NY 10065, USA
[5] Department of Genomics and Computational Biology, University of Massachusetts Chan Medical School, MA 01605, USA

**SUPPLEMENTARY TABLES:**

**Table S1:** List of cancer mutation studies, TFs, TF motifs, and chromatin accessibility data used in this study.

**Table S2:** Minimum enrichment q-values within the binding site for each <TF-cluster, tissue> pair.

**Table S3:** Percentage of mutations contributed by each COSMIC SBS signature for all mutations and enriched TF binding site mutations.

**Table S4:** Percentage of mutations contributed by each COSMIC SBS signature at each enriched TF cluster, per tissue.

#### SUPPLEMENTARY FIGURES

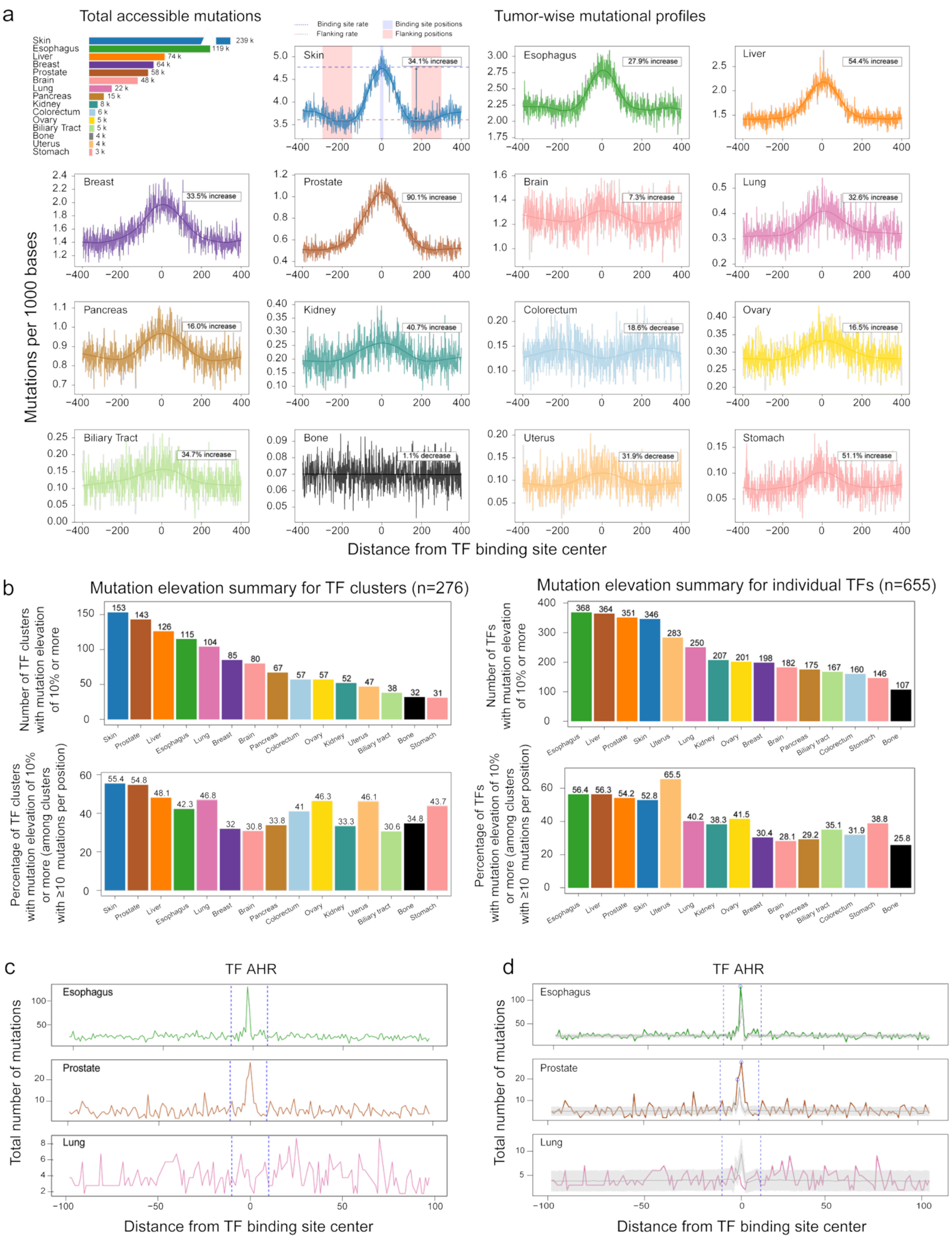


**Figure S1: Widespread elevation in mutation rates at TF binding sites.**

**(a)** Total number of accessible mutations for each tissue type analyzed, and tumor-wise aggregated mutational profiles at binding sites. Tumors are arranged in order of decreasing number of available mutations. Each profile is annotated with the elevation in mutation rate seen at binding site (+/-10bp from center) compared to the flanks (150-300bp from center), as described in Methods.

**(b)** Cluster-wise and TF-wise analysis of elevation at mutation rates within binding sites was done as described in Methods. On the left, results are shown at the level of TF clusters, with the number of clusters showing (more than 10%) elevation on the top, and the percentage of available TF clusters elevated (defined as those with more than 10 mutations in both flanks and binding site, Methods). On the right, same analysis is shown at the level of individual TFs (Methods).
**(c)** Example of distinct mutation profiles for a TF-cluster (AHR) in multiple tissue types (Esophagus, Prostate and Lung). Each profile shows observed mutation counts for each position (0-100 bp from binding site center) from all tumors in that tissue type. Blue dashed lines represent the binding site region (considered to be +/-10bp from the center).
**(d)** Same mutation profile as in (c), with the addition of expected mutation rates, shown as mean +/-2 standard deviations of the sequence-derived expected mutation counts (gray, Methods**)**

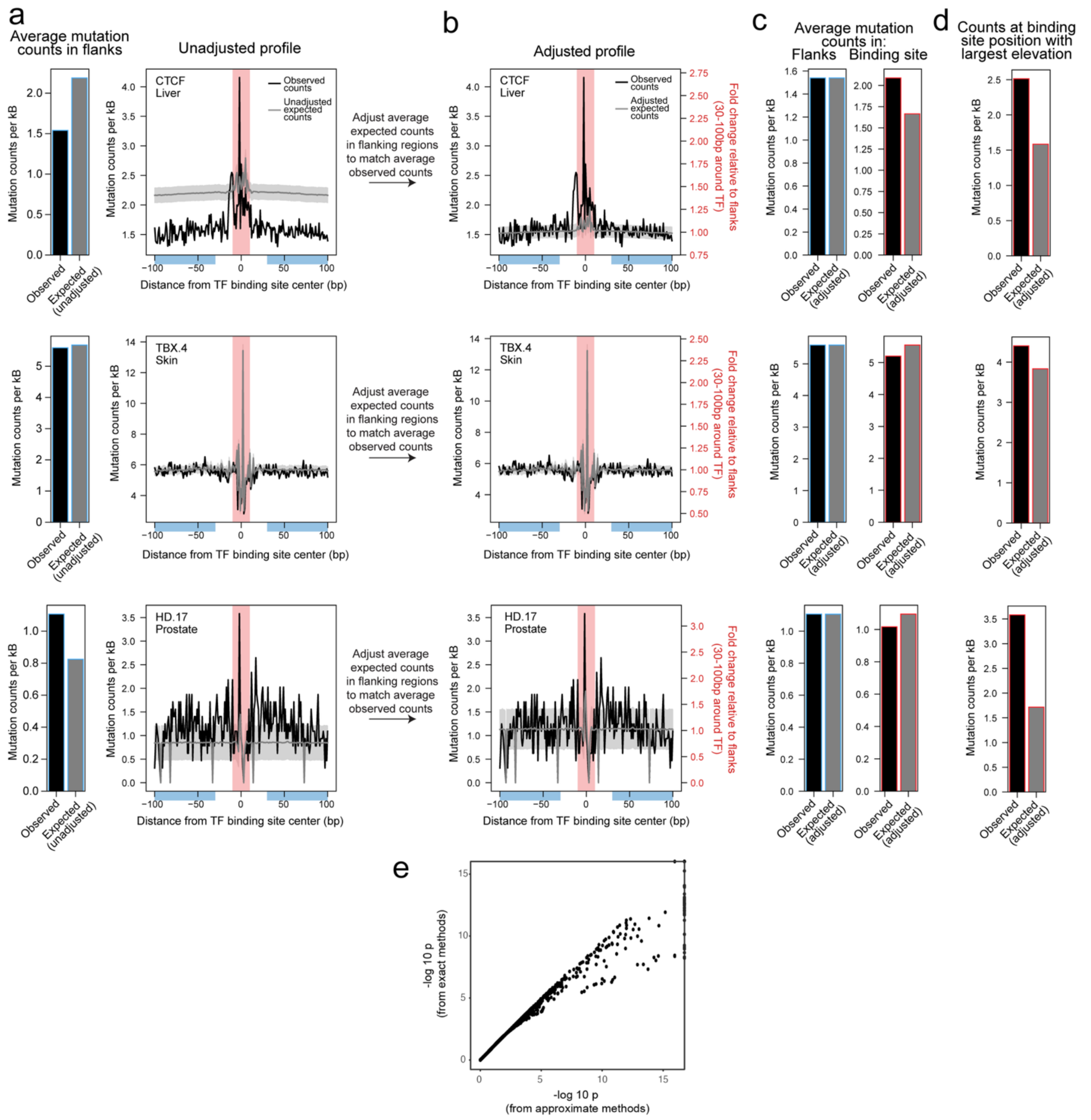


**Figure S2: Adjustment of expected mutation profiles to account for local variation in mutation rates and computation of p-values for enrichment tests.**

**(a)** Unadjusted profiles of mutation counts (gray) computed for TF binding sites and their flanking regions, by summing up expected counts at observed trimers at each position (**Methods**). In these unadjusted profiles, the rates observed in the local window differ between expected and observed counts, potentially due to local differences in mutation rates not associated with TF binding (e.g. due to cellular factors such as replication timing, transcription level, or chromatin environment). Three examples are shown, where the expected unadjusted counts are higher (top), similar (center), or lower (bottom) than the observed counts. The profiles for the observed counts are shown in black.

**(b)** Following similar approaches to existing work, we adjust expected trimer mutation rates so that the average expected mutation counts in flanking regions match the average observed counts (**Methods**).

**(c)** Left: after the normalization procedure, average observed mutation counts in the flanks match the average expected counts. Right: in the TF binding site regions (defined here at +/-10bp from the binding site center), the adjusted expected counts can differ from observed counts.

**(d)** At individual positions within the TF binding sites, observed mutation counts can differ from expected counts. Mutation counts shown are for the binding site positions with the largest elevation in actual counts compared to expected counts.

**(e)** Comparison of p-value computations for a subset of binding site positions computed using both the exact (DFT-CF) and the much faster, approximate (RNA) form of the Poisson binomial. Based on these comparisons, all positions were first computed using the RNA form, with exact forms then computed for the small number of positions with RNA p-values below 1e-4.

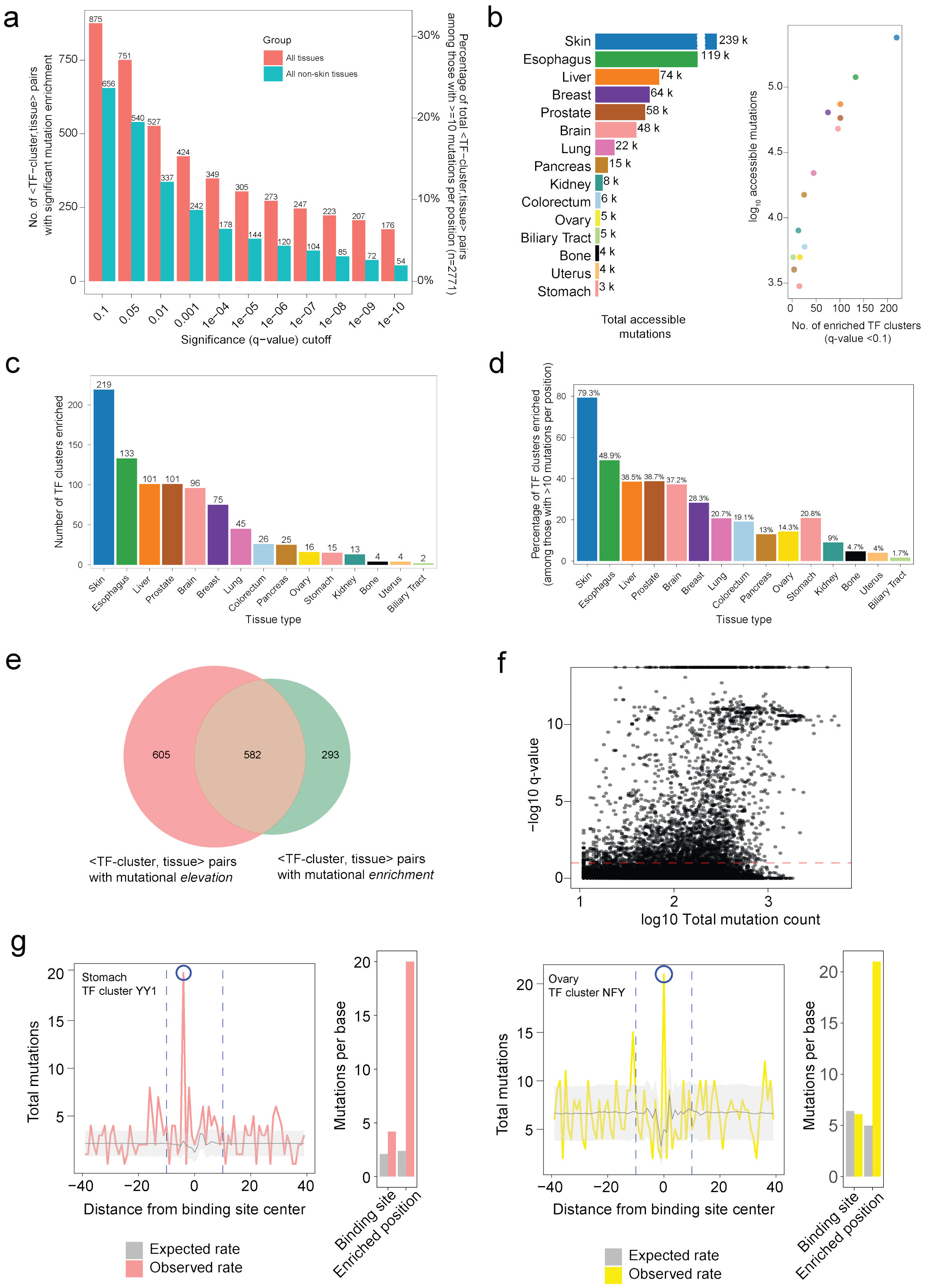


**Figure S3: Analysis of enriched <TF-cluster, tissue> pairs.**

**(a)** Number of <TF-cluster, tissue> pairs where at least one TF binding site position shows significant mutation enrichment, at different q-value cutoffs. Numbers are shown for all tissues (pink) and all tissues excluding skin (teal). Secondary axis represents these numbers as a percentage of total <TF-cluster, tissue> pairs with sufficient mutation counts, i.e. after filtering TFs 10 or fewer mutations at all binding site positions.

**(b)** Left: Total number of mutations in accessible regions of the genome for tumors from each tissue type. When both DNase and ATAC data were available (Methods), the intersection was used for these counts. For 3/15 tissues only DNase was available, and used for these counts. Right: Total number of enriched <TF-cluster, tissue> pairs for each tissue type, plotted against the (log) mutation count in accessible regions.
**(c)** Number and **(d)** Percentage of enriched <TF-cluster, tissue> pairs, at a q-value cutoff of 0.1. Percentage numbers are within <TF-cluster, tissue> pairs with sufficient mutation counts for analysis (as in (a)).
**(e)** Overlap between <TF-cluster, tissue> pairs identified as having an overall elevation in mutation rates at binding sites (see Fig. S1) and <TF-cluster, tissue> identified as having a potential enrichment over expected mutation rates (q-value <0.1) following our workflow.
**(f)** Observed mutation count at each binding site position (log10, x-axis) and the significance of enrichment over expected mutation count (-log10 q-value, y-axis). Red line represents a q-value cutoff of 0.1, above which a position might be considered significantly enriched. Only binding site positions with more than 10 observed mutation counts are shown, across all TF and tissue combinations analyzed.
**(g)** Examples of mutation profiles (left) for TFs where specific binding site positions show significant enrichment (q-value below 0.1), with a substantially higher increase in mutation rate over the expected rate (gray) than when looking at the entire binding site as a whole (right).

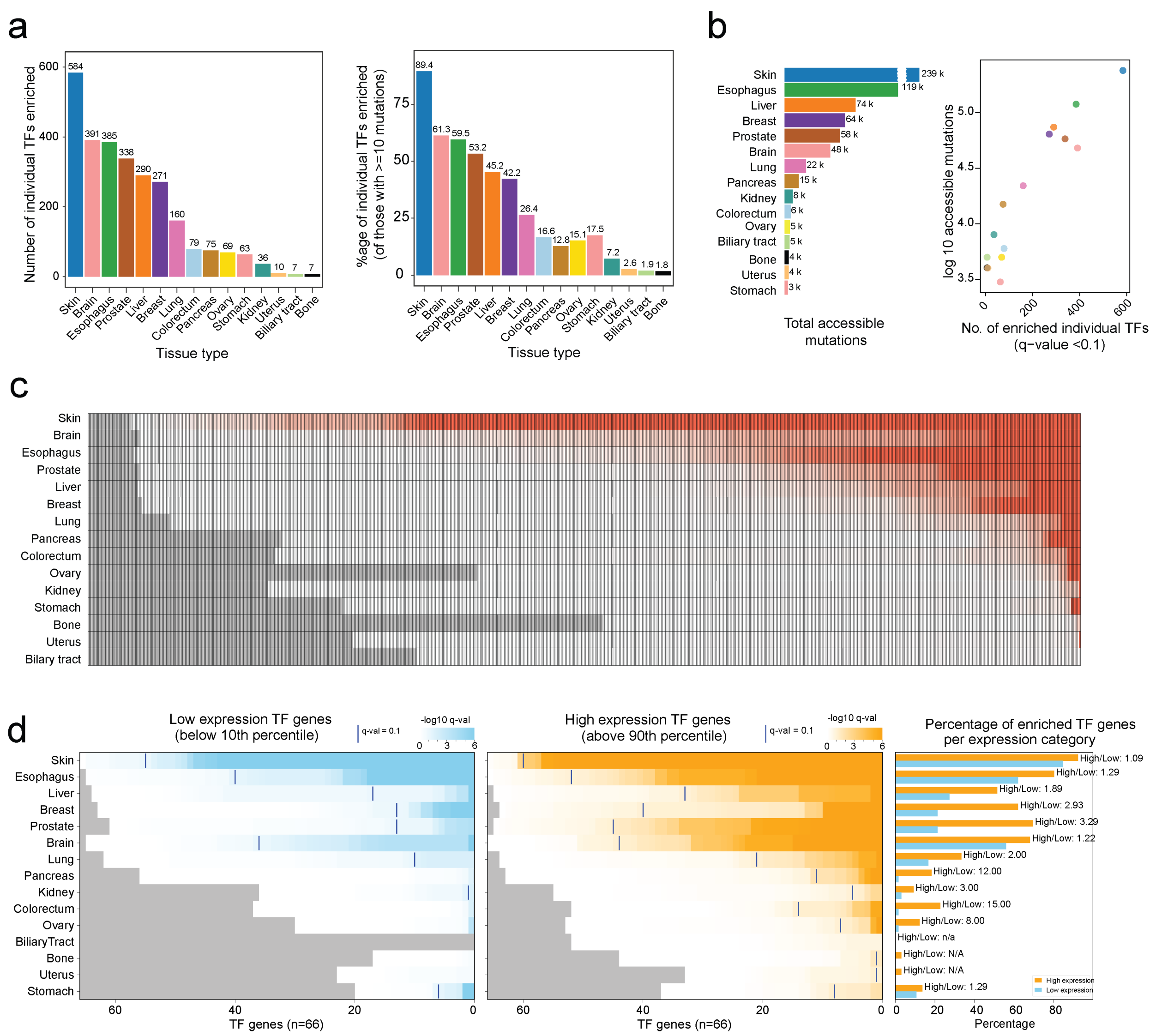


**Figure S4: Analysis of enriched <TF, tissue> pairs.**

**(a)** Left: Number and Right**:** Percentage of enriched < TF, tissue > pairs, at a q-value cutoff of 0.1. Percentage numbers are within < TF, tissue > pairs with sufficient mutation counts for analysis (as in **Fig. S3**).

**(b)** Same as **Fig. S3b**, repeated for individual enriched TF counts instead of TF-clusters.

**(c)** Same heatmap as **Fig. 2a**, repeated for individual TFs (n=655) instead of TF clusters.

**(d)** Heatmap for highly expressed TFs as in **Fig. 2d**, repeated for TFs below 10th percentile counted in the low expression group and TFs above 90th percentile counted in the high expression

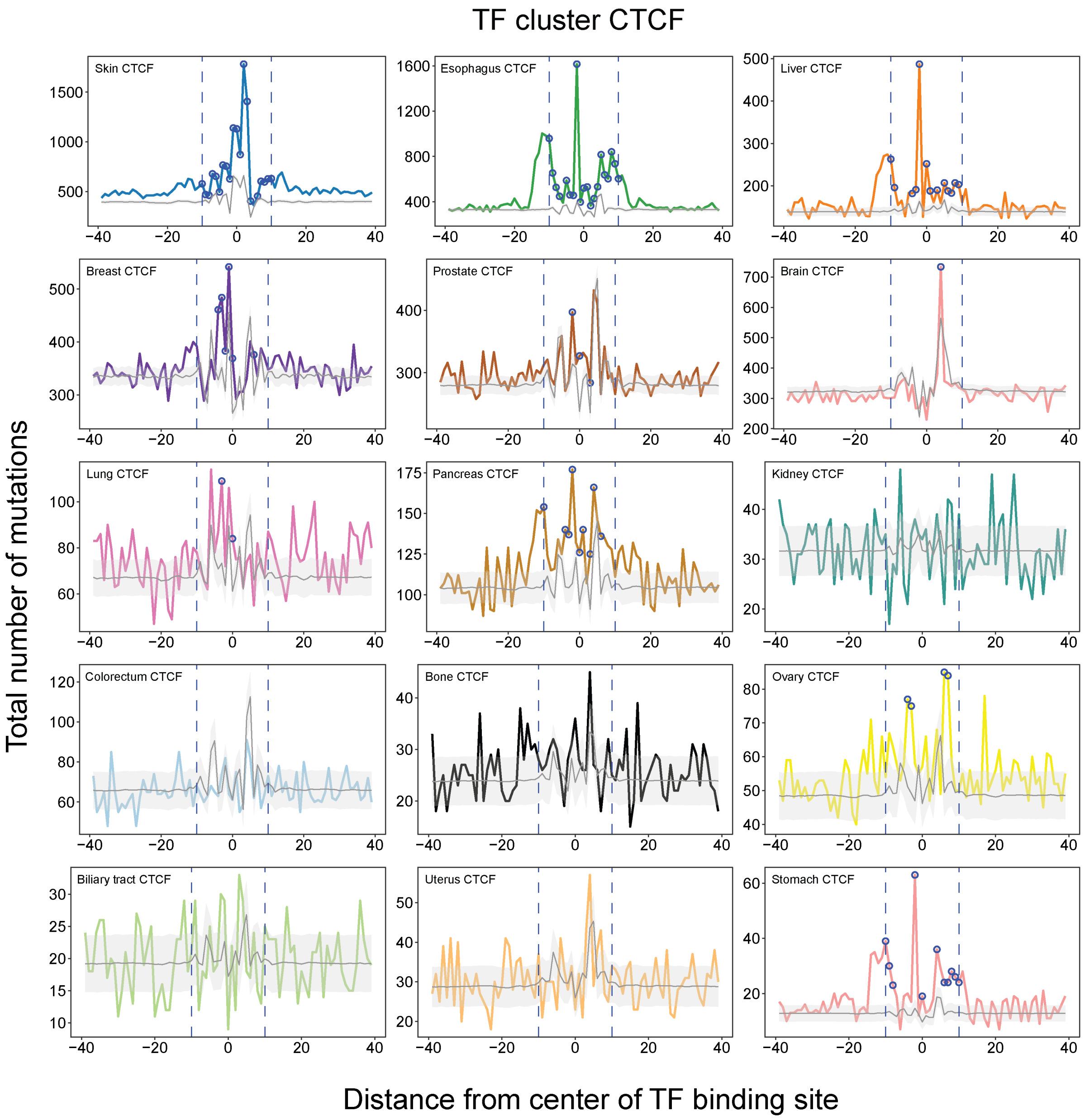


**Figure S5: Mutational profiles for TF cluster CTCF, in all tissue types analyzed.** Mutational profiles are drawn similar to **Fig. 3a**, for all tissue types included. Different positions within the binding site enriched in each tissue (when present) are highlighted.

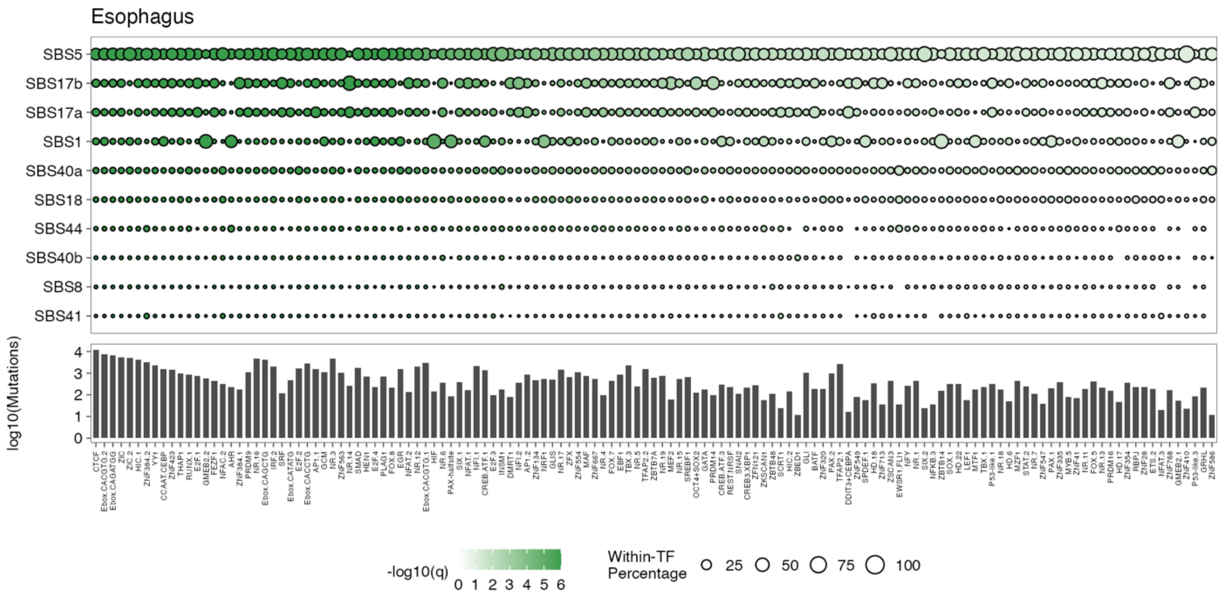


**Figure S6: SBS signatures contributing to enriched-position mutations, for each enriched TF cluster in esophageal tumors.**

As **Fig. 4c**, except all esophageal enriched TFs are shown instead of top 75. Signature rows are the top 10 signatures with most attributed mutations at enriched binding site positions.

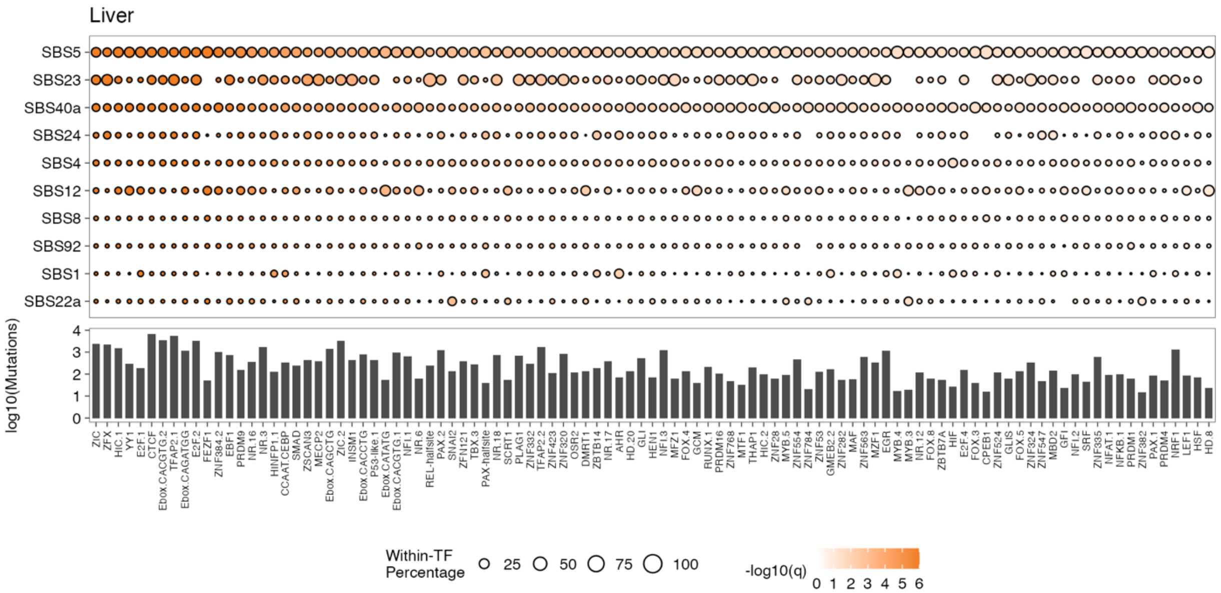


**Figure S7: SBS signatures contributing to enriched-position mutations, for each enriched TF cluster in liver tumors.**

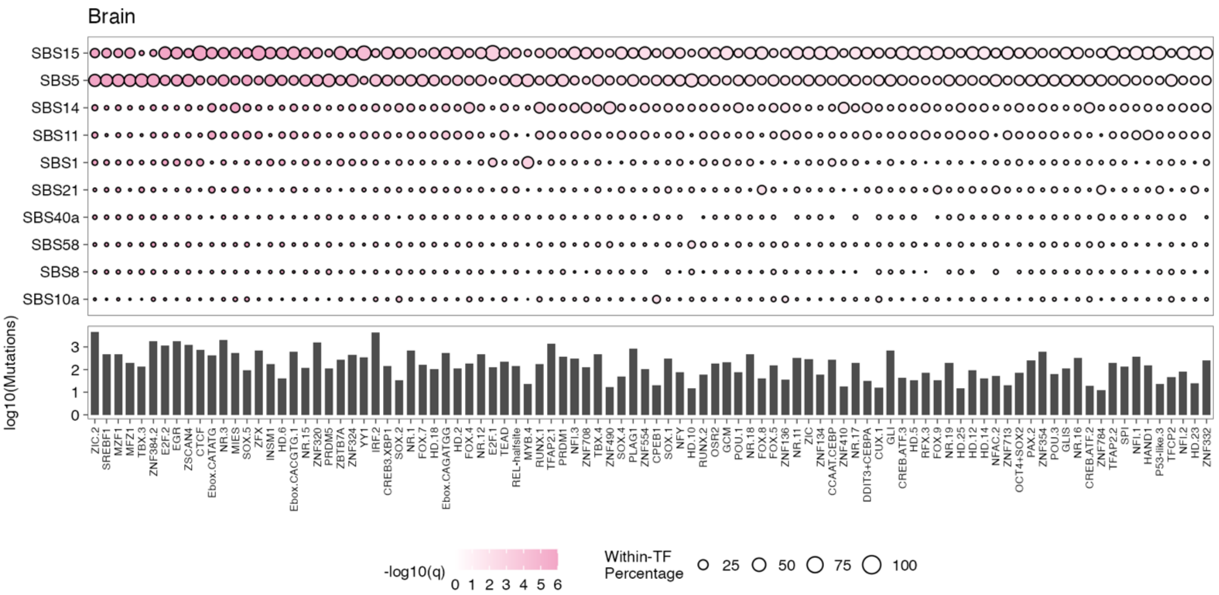


**Figure S8: SBS signatures contributing to enriched-position mutations, for each enriched TF cluster in brain tumors.**

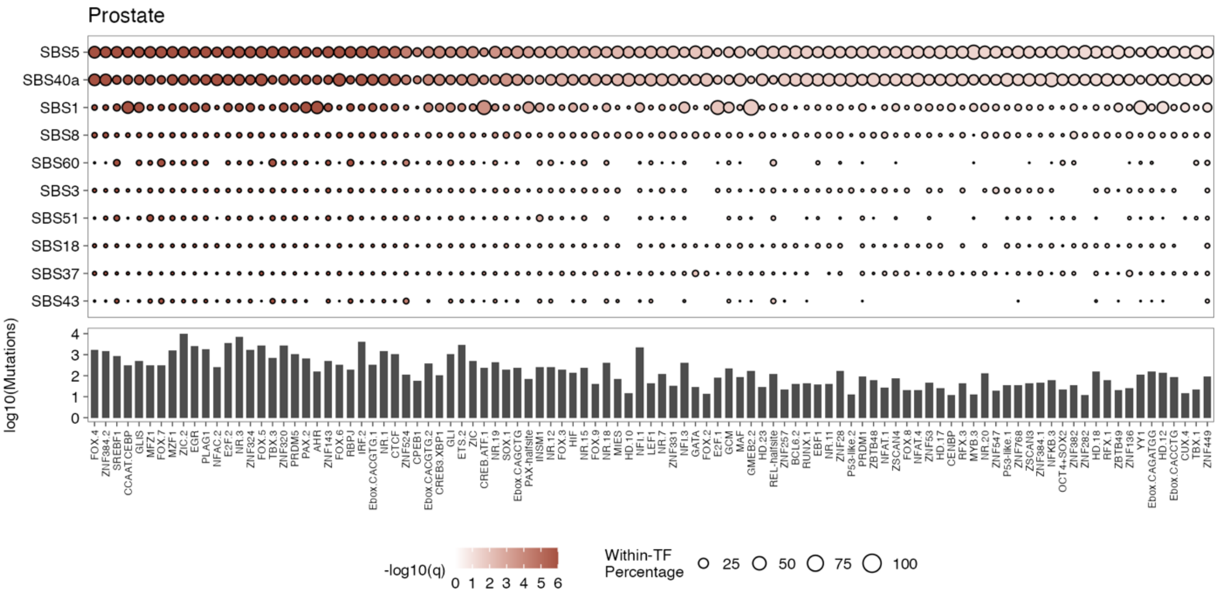


**Figure S9: SBS signatures contributing to enriched-position mutations, for each enriched TF cluster in prostate tumors.**

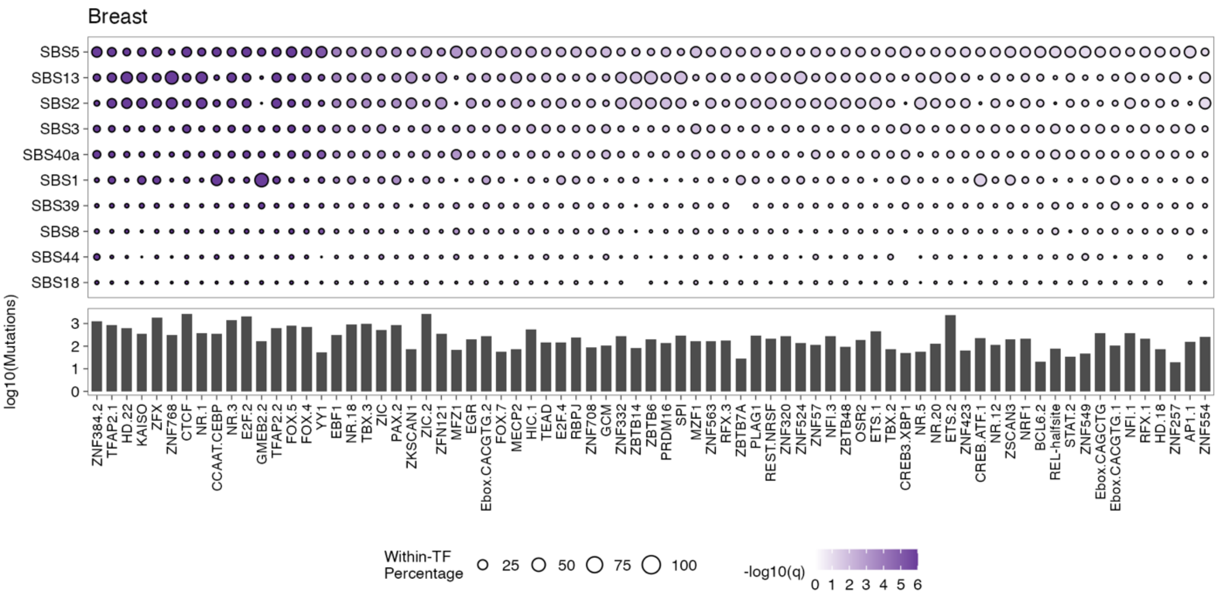


**Figure S10: SBS signatures contributing to enriched-position mutations, for each enriched TF cluster in breast tumors.**

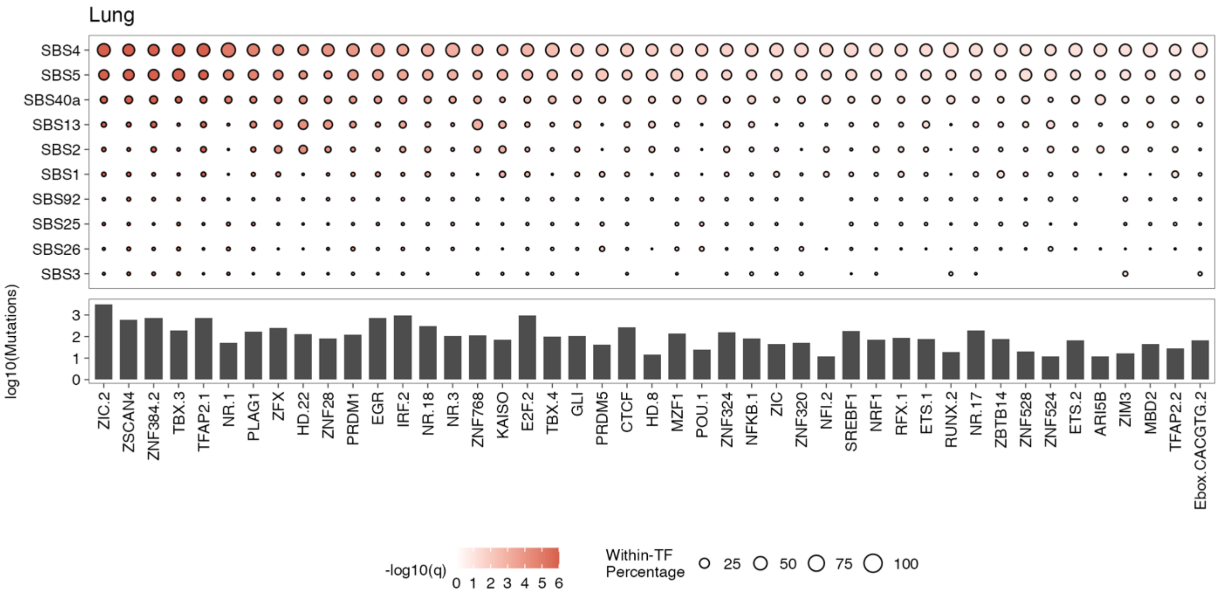


**Figure S11: SBS signatures contributing to enriched-position mutations, for each enriched TF cluster in lung tumors.**

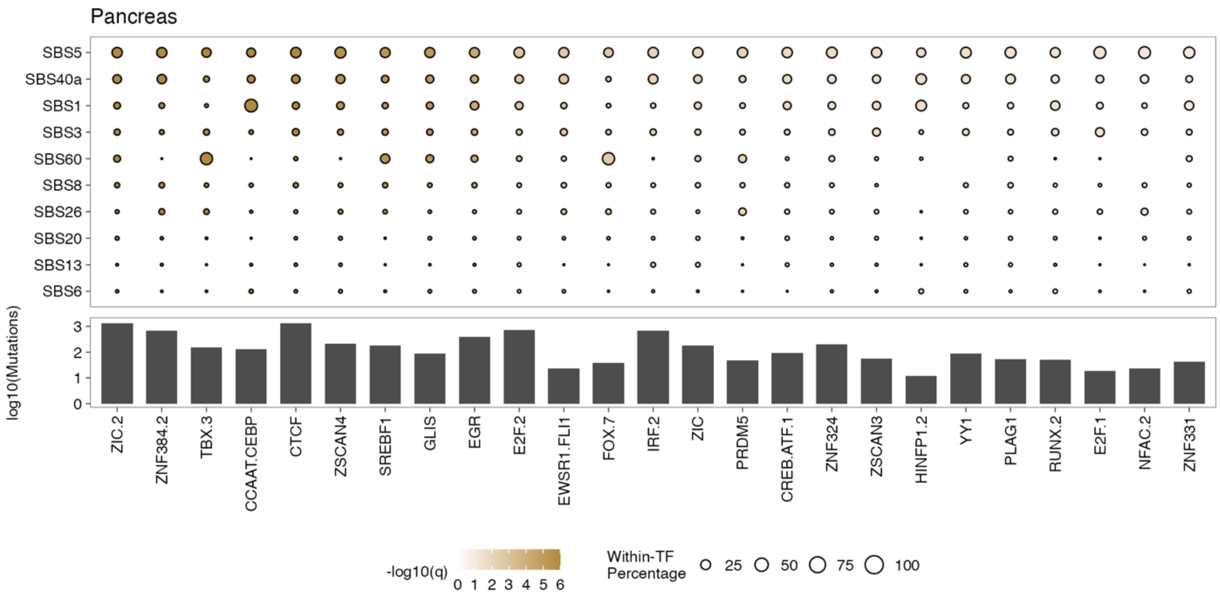


**Figure S12: SBS signatures contributing to enriched-position mutations, for each enriched TF cluster in pancreatic tumors.**

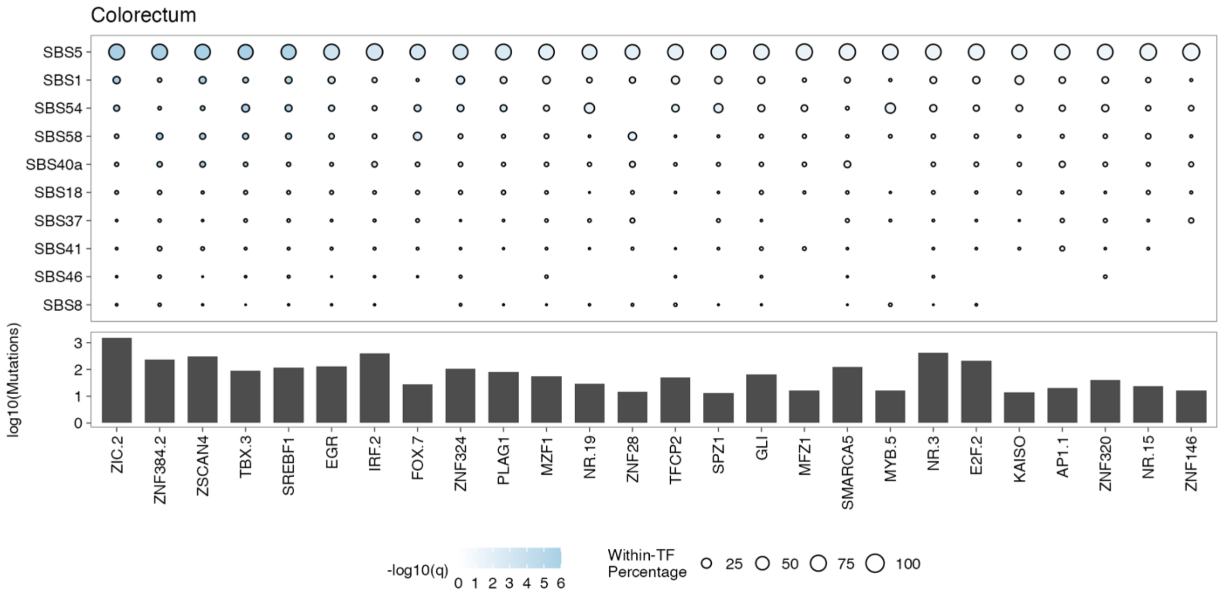


**Figure S13: SBS signatures contributing to enriched-position mutations, for each enriched TF cluster in colorectal tumors.**

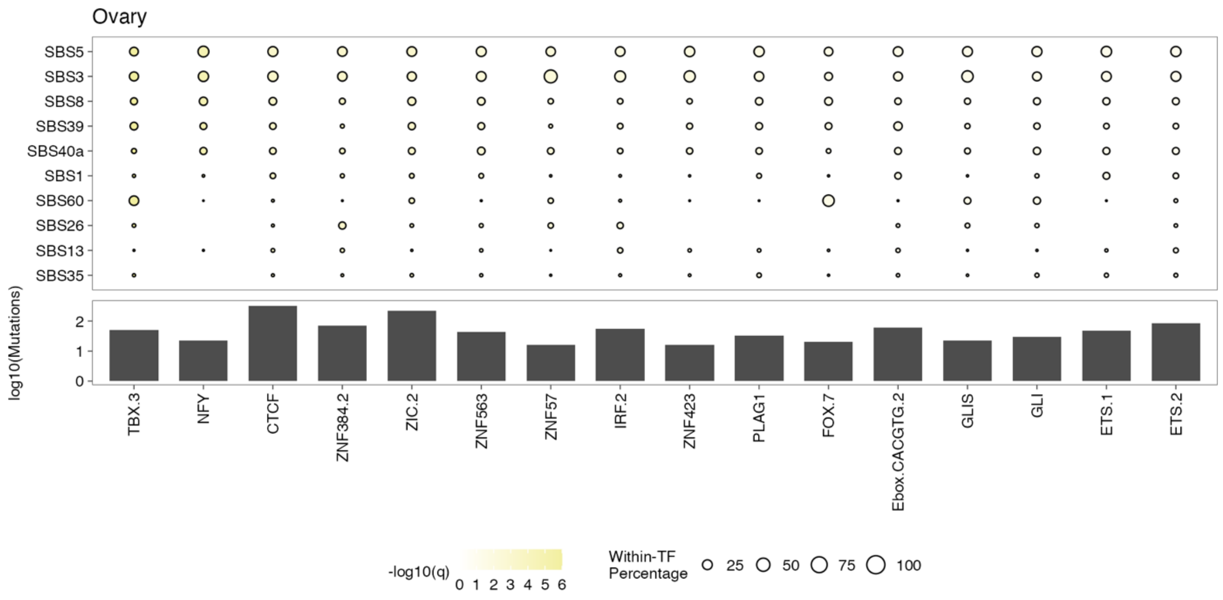


**Figure S14: SBS signatures contributing to enriched-position mutations, for each enriched TF cluster in ovarian tumors.**

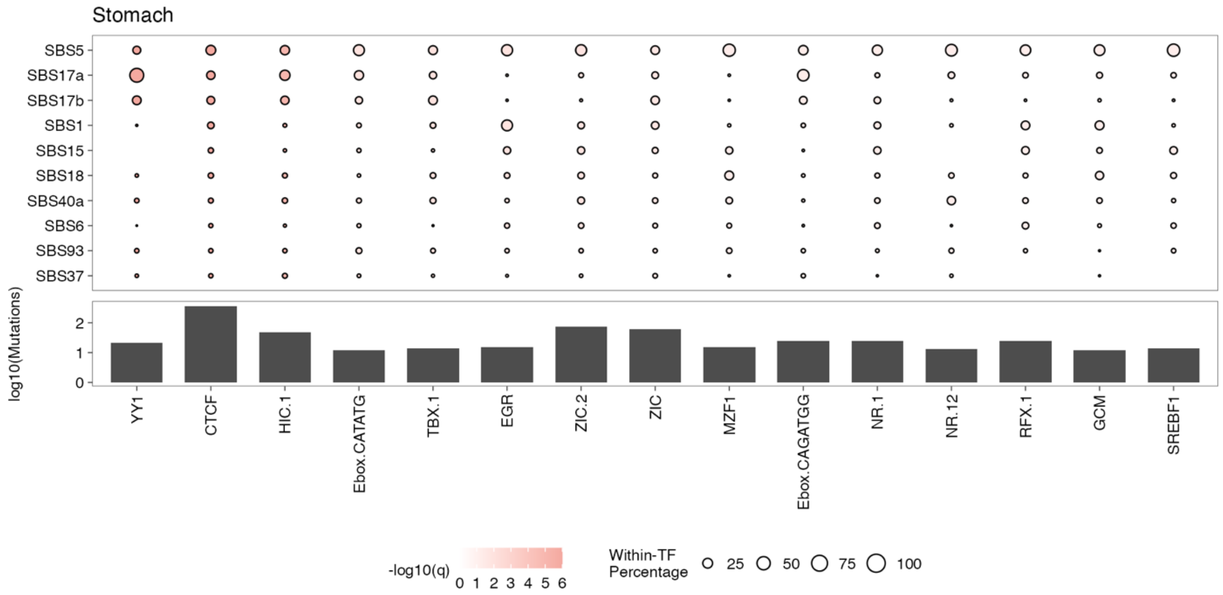


**Figure S15: SBS signatures contributing to enriched-position mutations, for each enriched TF cluster in stomach tumors.**

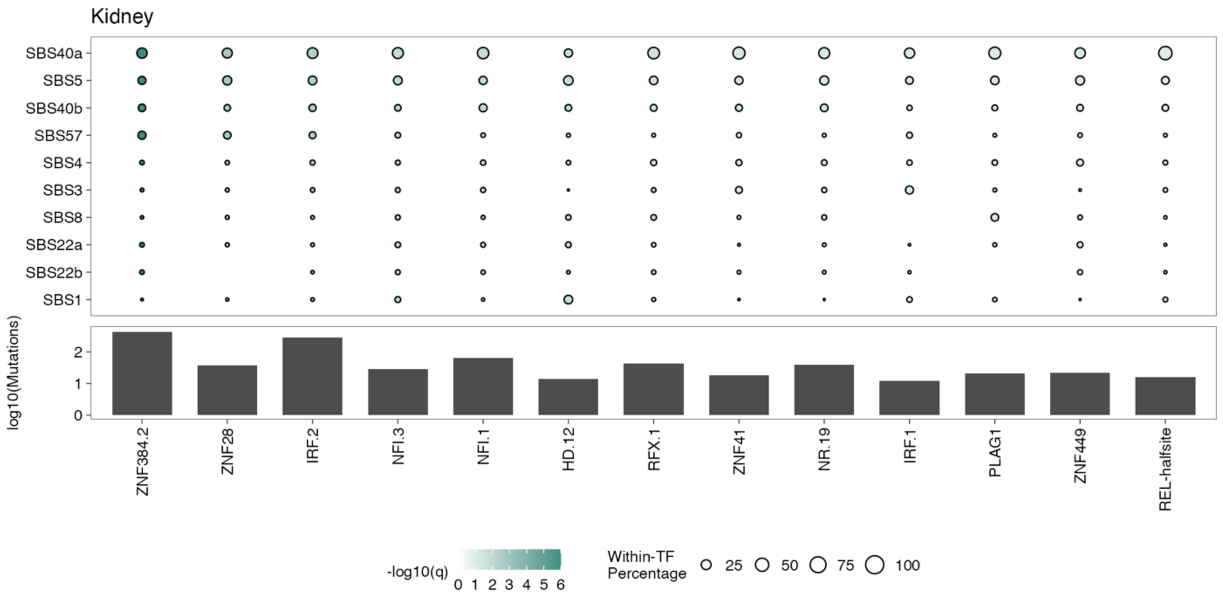


**Figure S16: SBS signatures contributing to enriched-position mutations, for each enriched TF cluster in kidney tumors.**
